# A genome-wide algal mutant library reveals a global view of genes required for eukaryotic photosynthesis

**DOI:** 10.1101/464859

**Authors:** Xiaobo Li, Weronika Patena, Friedrich Fauser, Robert E. Jinkerson, Shai Saroussi, Moritz T. Meyer, Nina Ivanova, Jacob M. Robertson, Rebecca Yue, Ru Zhang, Josep Vilarrasa-Blasi, Tyler M. Wittkopp, Silvia Ramundo, Sean R. Blum, Audrey Goh, Matthew Laudon, Tharan Srikumar, Paul A. Lefebvre, Arthur R. Grossman, Martin C. Jonikas

**Author notes:** Present addresses: School of Life Sciences, Westlake University, Hangzhou, Zhejiang Province 310064, China (X.L.); Department of Chemical and Environmental Engineering, University of California, Riverside, California 92521, USA (R.E.J); Donald Danforth Plant Science Center, 975 North Warson Road, St. Louis, Missouri 63132, USA (R.Z.); Salk Institute for Biological Studies, La Jolla, California 92037, USA (T.M.W.).

## Abstract

Photosynthetic organisms provide food and energy for nearly all life on Earth, yet half of their protein-coding genes remain uncharacterized^1,2^. Characterization of these genes could be greatly accelerated by new genetic resources for unicellular organisms that complement the use of multicellular plants by enabling higher-throughput studies. Here, we generated a genome-wide, indexed library of mapped insertion mutants for the flagship unicellular alga *Chlamydomonas reinhardtii* (Chlamydomonas hereafter). The 62,389 mutants in the library, covering 83% of nuclear, protein-coding genes, are available to the community. Each mutant contains unique DNA barcodes, allowing the collection to be screened as a pool. We leveraged this feature to perform a genome-wide survey of genes required for photosynthesis, which identified 303 candidate genes. Characterization of one of these genes, the conserved predicted phosphatase *CPL3*, showed it is important for accumulation of multiple photosynthetic protein complexes. Strikingly, 21 of the 43 highest-confidence genes are novel, opening new opportunities for advances in our understanding of this biogeochemically fundamental process. This library is the first genome-wide mapped mutant resource in any unicellular photosynthetic organism, and will accelerate the characterization of thousands of genes in algae, plants and animals.

---

Among unicellular photosynthetic organisms, the green alga Chlamydomonas has long been employed for genetic studies of eukaryotic photosynthesis because of its rare ability to grow in the absence of photosynthetic function^3^. In addition, it has made extensive contributions to our basic understanding of light signaling, stress acclimation, and metabolism of carbohydrates, lipids, and pigments (Fig. 1a)^4-6^. Moreover, Chlamydomonas retained many genes from the plant-animal common ancestor, which allowed it to reveal fundamental aspects of the structure and function of cilia and basal bodies^7,8^. Like *Saccharomyces cerevisiae*, Chlamydomonas can grow as a haploid, facilitating genetic studies. However, until now, the value of Chlamydomonas has been limited by the lack of mutants in most of its nuclear genes.

**Fig. 1.**
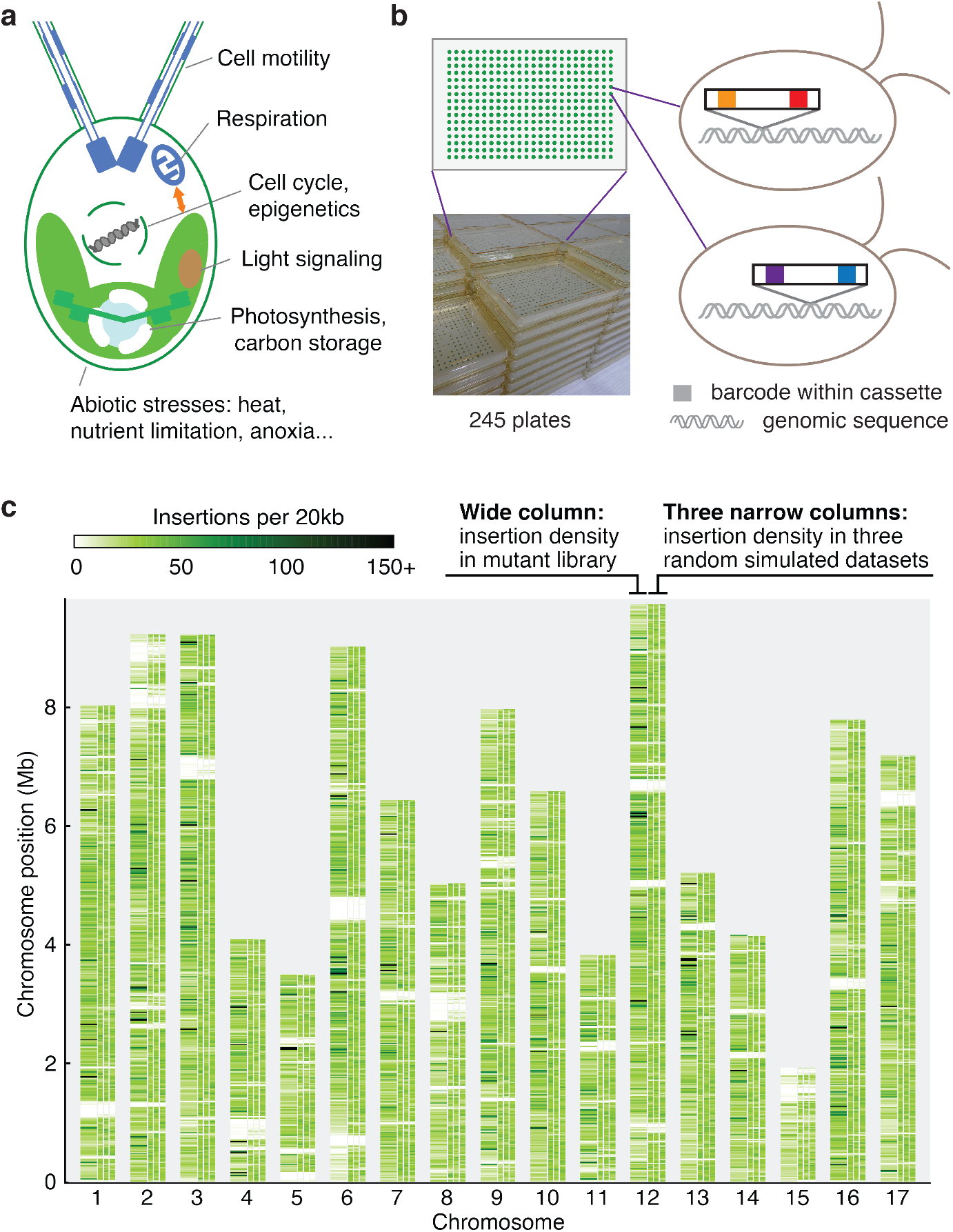
A genome-wide library of Chlamydomonas mutants was generated by random insertion of barcoded cassettes and mapping of insertion sites. **a**, *Chlamydomonas reinhardtii* is used for studies of various cellular processes and organism-environment interactions. **b**, Our library contains 62,389 insertional mutants maintained as 245 plates of 384-colony arrays. Each mutant contains at least one insertion cassette at a random site in its genome; each insertion cassette contains one unique barcode at each end (Supplementary Fig. 1a-c). **c**, The insertion density is largely random over the majority of the genome. This panel compares the observed insertion density over the genome (the left column above each chromosome number) to three simulations with insertions randomly distributed over all mappable positions in the genome (the three narrow columns to the right for each chromosome). Areas that are white throughout all columns represent regions where insertions cannot be mapped to a unique genomic position due to highly repetitive sequence. See also Supplementary Fig. 4.

In the present study, we sought to generate a genome-wide collection of Chlamydomonas mutants with known gene disruptions to provide mutants in genes of interest for the scientific community, and then to leverage this collection to reveal genes with roles in photosynthesis. To reach the necessary scale, we chose to use random insertional mutagenesis and built on advances in insertion mapping and mutant propagation from our pilot study^9^. To enable mapping of insertion sites and screening pools of mutants on a much larger scale, we developed new tools leveraging unique DNA barcodes in each transforming cassette.

We generated mutants by transforming haploid cells with DNA cassettes that randomly insert into the genome and inactivate the genes they insert into. We maintained the mutants as indexed colony arrays on agar media containing acetate as a carbon and energy source to allow recovery of mutants with defects in photosynthesis. Each DNA cassette contained two unique barcodes, one on each side of the cassette (Supplementary Fig. 1a-d). For each mutant, the barcode and genomic flanking sequences on each side of the cassette were initially unknown (Supplementary Fig. 1e). We determined the sequence of the barcode(s) in each mutant colony by combinatorial pooling and deep sequencing (Supplementary Fig. 1f). We then mapped each insertion by pooling all mutants and amplifying all flanking sequences together with their corresponding barcodes followed by deep sequencing (Supplementary Fig. 1g). The combination of these datasets revealed the insertion site(s) in each mutant. This procedure yielded 62,389 mutants on 245 plates, with a total of 74,923 insertions that were largely randomly distributed over the chromosomes (Fig. 1, b and c, and Supplementary Table 5).

This library provides mutants for ~83% of all nuclear genes (Fig. 2a-d). Approximately 69% of genes are represented by an insertion in a 5’ UTR, an exon or an intron – regions most likely to cause an altered phenotype when disrupted. Many gene sets of interest to the research community are well represented, including genes encoding proteins phylogenetically associated with the plant lineage (GreenCut2)^1^, proteins that localize to the chloroplast^10^, or those associated with the structure and function of flagella or basal bodies^11,12^ (Fig. 2b). Mutants in this collection are available through the website https://www.chlamylibrary.org/. Over 1,800 mutants have already been distributed to over 200 laboratories worldwide in the first 18 months of pre-publication distribution (Fig. 2e). These mutants are facilitating genetic investigation of a broad range of processes, ranging from photosynthesis and metabolism to cilia structure and function (Fig. 2f).

**Fig. 2.**
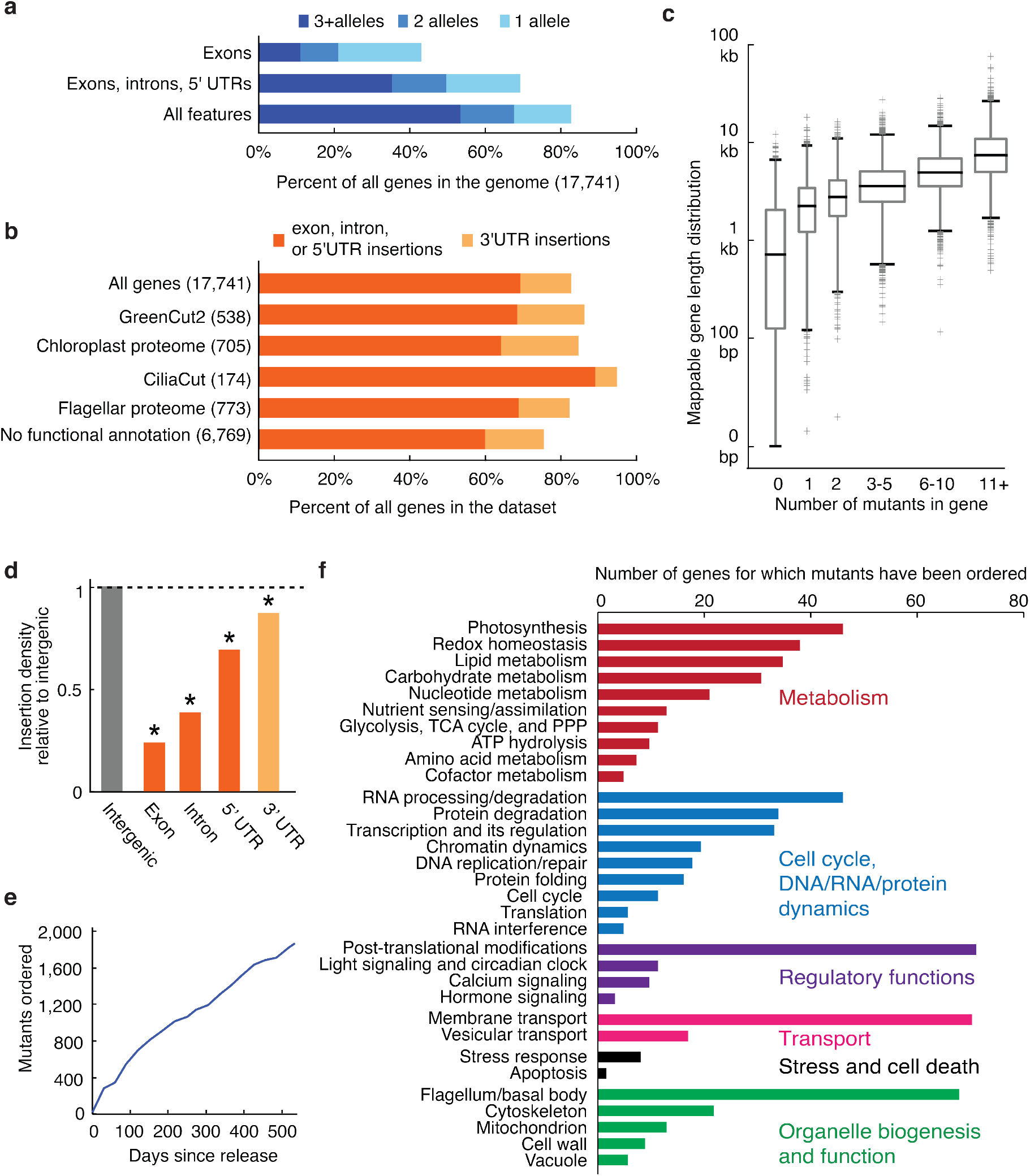
The library covers 83% of Chlamydomonas genes. **a**, 83% of all Chlamydomonas genes have one or more insertions in the library. **b**, In various functional groups, more than 75% of genes are represented by insertions in the library. **c**, The number of insertions per gene is roughly correlated with gene length. Box heights represent quartiles, whiskers represent 1^st^ and 99^th^ percentiles, and outliers are plotted as crosses. Box widths are proportional to the number of genes in each bin. **d**, Insertion density varies among different gene features, with the lowest density in exons. **e**, More than 1,800 mutants were distributed to approximately 200 laboratories in the world during the first 18 months of its availability. **f**, Distributed mutants are being used to study a variety of biological processes. Only genes with some functional annotation are shown.

To identify genes required for photosynthesis, we screened our library for mutants deficient in photosynthetic growth. Rather than phenotyping each strain individually, we pooled the entire library into one culture and leveraged the unique barcodes present in each strain to track its abundance after growth under different conditions. This feature enables genome-wide screens with speed and depth unprecedented in photosynthetic eukaryotes. We grew a pool of mutants photosynthetically in light in minimal Tris-Phosphate (TP) medium with CO_2_ as the sole source of carbon, and heterotrophically in the dark in Tris-Acetate-Phosphate (TAP) medium, where acetate provides fixed carbon and energy^3^ (Fig. 3a). To quantify mutant growth under each condition, we amplified and deep sequenced the barcodes from the final cell populations. We then compared the ability of each mutant to grow under photosynthetic and heterotrophic conditions by comparing the read counts of each barcode from each condition (Supplementary Table 10; Methods). Mutant phenotypes were highly reproducible (Fig. 3b and Supplementary Fig. 5, a and b). We identified 3,109 mutants deficient in photosynthetic growth (Fig. 3c and Methods).

**Fig. 3.**
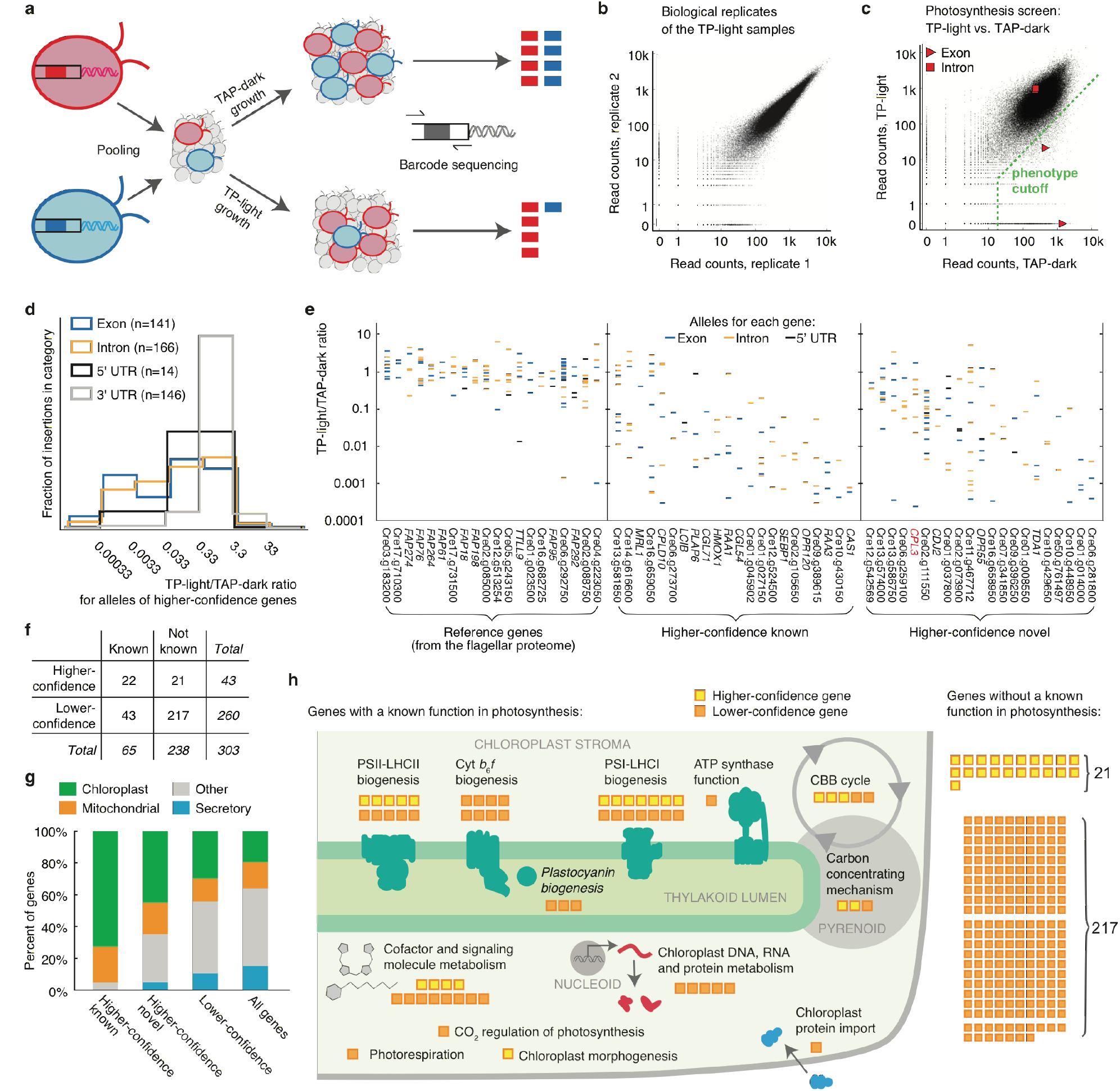
A high-throughput screen using the library identifies many genes with known roles in photosynthesis and reveals many novel components. **a**, Unique barcodes allow screening mutants in a pool. Mutants deficient in photosynthesis can be identified because their barcodes will be less abundant after growth in photosynthetic (TP-light) relative to after growth in heterotrophic (TAP-dark) conditions. **b**, Biological replicates were highly reproducible, with a Spearman’s correlation of 0.982. Each dot represents one barcode. See also Supplementary Fig. 5 and Methods. **c**, The phenotype of each insertion was determined by comparing its read count under TAP-dark and TP-light conditions. Insertions that fell below the phenotype cutoff were considered to show a defect in photosynthesis. cpl3 alleles are highlighted in red squares or triangles. **d**, Insertion phenotypes vary depending on the gene feature disrupted: exon and intron insertions are most likely to show strong phenotypes, while 3’UTR insertions rarely do. The plot is based on all insertions for the 43 higher-confidence genes. **e**, The TP-light/TAP-dark ratio of all the alleles are shown for hit and control genes. Each column is a gene; each horizontal bar is an allele, color-coded by feature. **f**, The 303 candidate genes were categorized according to (1) whether or not they were previously known to play a role in photosynthesis, and (2) whether the screen data yielded higher or lower confidence that mutation of that gene causes a defect in photosynthetic growth. g, Known higher-confidence genes, novel higher-confidence genes, and lower-confidence genes are all enriched in predicted chloroplast-targeted proteins (P h, Twenty-two of the higher-confidence genes and 43 of the lower-confidence genes were previously known to have a role in processes related to photosynthesis. The screen additionally identified 21 higher-confidence and 217 lower-confidence genes that were not previously known to be involved in photosynthesis.

To identify genes with roles in photosynthesis, we developed a statistical analysis framework that leverages the presence of multiple alleles for many genes. This framework allows us to overcome several sources of false positives that have been difficult to identify with previous methods, including cases where the phenotype is not caused by the mapped disruption. For each gene, we counted the number of mutant alleles with and without a phenotype, and evaluated the likelihood of obtaining these numbers by chance given the total number of mutants in the library that exhibit the phenotype (Supplementary Table 11; Methods).

We identified 303 candidate photosynthesis genes based on our statistical analysis above. These genes are enriched for membership in a diurnally regulated photosynthesis-related transcriptional cluster^13^ (*P*<10^-11^), are enriched for upregulation upon dark-to-light transitions^14^ (*P*<0.003), and encode proteins enriched for predicted chloroplast localization (*P*<10^-8^). As expected^15^, the candidate genes also encode a disproportionate number of GreenCut2 proteins (*P*<10^-8^), which are conserved among photosynthetic organisms but absent from non-photosynthetic organisms^1^: 32 GreenCut2 proteins are encoded by the 303 candidate genes (11%), compared to |3% in the entire genome.

Photosynthesis occurs in two stages: the light reactions and carbon fixation. The light reactions convert solar energy into chemical energy, and require coordinated action of Photosystem II (PSII), Cytochrome *b*_6_*f*, Photosystem I (PSI), ATP synthase complexes, a plastocyanin or cytochrome *c*_6_ metalloprotein, as well as small molecule cofactors^16^. PSII and PSI are each assisted by peripheral light-harvesting complexes (LHCs) known as LHCII and LHCI, respectively. Carbon fixation is performed by enzymes in the Calvin-Benson-Bassham cycle, including the CO_2_-fixing enzyme Rubisco. In addition, most eukaryotic algae have a mechanism to concentrate CO_2_ around Rubisco to enhance its activity^17^.

Sixty-five of the genes we identified encode proteins that were previously shown to play a role in photosynthesis or chloroplast function in Chlamydomonas or vascular plants (Fig. 3f). These include three PSII-LHCII subunits (PSBP1, PSBP2, and PSB27) and seven PSII-LHCII biogenesis factors (CGL54, CPLD10, HCF136, LPA1, MBB1, TBC2, and Cre02.g105650), two cytochrome *b*_6_*f* complex subunits (PETC and PETM) and six cytochrome *b*_6_*f* biogenesis factors (CCB2, CCS5, CPLD43, CPLD49, MCD1, and MCG1), five PSI-LHCI subunits (LHCA3, LHCA7, PSAD, PSAE, and PSAL) and nine PSI-LHCI biogenesis factors (CGL71, CPLD46, OPR120, RAA1, RAA2, RAA3, RAT2, Cre01.g045902, and Cre09.g389615), one protein required for ATP synthase function (PHT3), plastocyanin (PCY1) and two plastocyanin biogenesis factors (CTP2 and PCC1), 12 proteins involved in the metabolism of photosynthesis cofactors or signaling molecules (CHLD, CTH1, CYP745A1, DVR1, HMOX1, HPD2, MTF1, PLAP6, UROD3, Cre08.g358538, Cre13.g581850, and Cre16.g659050), three Calvin-Benson-Bassham Cycle enzymes (FBP1, PRK1, and SEBP1), two Rubisco biogenesis factors (MRL1 and RMT2), three proteins involved in the algal carbon concentrating mechanism (CAH3, CAS1, and LCIB), as well as proteins that play a role in photorespiration (GSF1), CO_2_ regulation of photosynthesis (Cre02.g146851), chloroplast morphogenesis (Cre14.g616600), chloroplast protein import (SDR17), and chloroplast DNA, RNA, and protein metabolism (DEG9, MSH1, MSRA1, TSM2, and Cre01.g010864) (Fig. 3h and Supplementary Table 12). We caution that not all genes previously demonstrated to be required for photosynthetic growth are detectable by this approach, especially the ones with paralogous genes in the genome, such as *RBCS1* and *RBCS2* that encode the small subunit of Rubisco^18^. Nonetheless, the large number of known factors recovered in our screen is a testament to the power of this approach.

In addition to recovering these 65 genes with known roles in photosynthesis, our analysis revealed 238 candidate genes with no previously reported role in photosynthesis (Methods). These 238 genes represent a rich set of targets to better understand photosynthesis. Because our screen likely yielded some false positives, we divided all genes into “higher-confidence” (*P*<0.0011; FDR< 0.27) and “lower-confidence” genes based on the number of alleles that supported each gene’s involvement in photosynthesis (Fig. 3d-f; Tables 1 and 2; Methods). The 21 higher-confidence genes with no previously reported role in photosynthesis are enriched in chloroplast localization (9/21, *P*<0.011; Fig. 3g) and transcriptional upregulation during dark to light transition (5/21, *P*<0.005), similar to the known photosynthesis genes. Thus, these 21 higher-confidence genes are particularly high-priority targets for the field to pursue.

Functional annotations for 15 of the 21 higher-confidence genes suggest that these genes could play roles in regulation of photosynthesis, photosynthetic metabolism, and biosynthesis of the photosynthetic machinery. Seven of the genes likely play roles in regulation of photosynthesis: *GEF1* encodes a voltage-gated channel, Cre01.g008550 and Cre02.g111550 encode putative protein kinases, *CPL3* encodes a predicted protein phosphatase, *TRX21* contains a thioredoxin domain, Cre12.g542569 encodes a putative glutamate receptor, and Cre13.g586750 contains a predicted nuclear importin domain. Six of the genes are likely involved in photosynthetic metabolism: the Arabidopsis homolog of Cre10.g448950 modulates sucrose and starch accumulation^19^, Cre11.g467712 contains a starch-binding domain, Cre02.g073900 encodes a putative carotenoid dioxygenase, *VTE5* encodes a putative phosphatidate cytidylyltransferase, Cre10.g429650 encodes a putative alpha/beta hydrolase, and Cre50.g761497 contains a magnesium transporter domain. Finally, two of the genes are likely to play roles in the biogenesis and function of photosynthesis machinery: *EIF2* has a translation initiation factor domain, and *CDJ2* has a chloroplast DnaJ domain. Future characterization of these genes by the community is likely to yield fundamental insights into our understanding of photosynthesis.

As an illustration of the value of genes identified in this screen, we sought to explore the specific function of one of the novel higher-confidence hits, *CPL3* (*Conserved in Plant Lineage 3*, Cre03.g185200), which encodes a putative protein phosphatase (Fig. 4a and Supplementary Fig. 6e). Many proteins in the photosynthetic apparatus are phosphorylated, but the role and regulation of these phosphorylations are poorly understood^20^. In our screen, three mutants in *CPL3* exhibited a deficiency in photosynthetic growth (Fig. 3c and Supplementary Table 13). We chose to examine one allele (LMJ.RY0402.153647, referred to hereafter as *cpl3*; Fig. 4a and Supplementary Fig. 6a) for phenotypic confirmation, genetic complementation, and further studies.

**Fig. 4.**
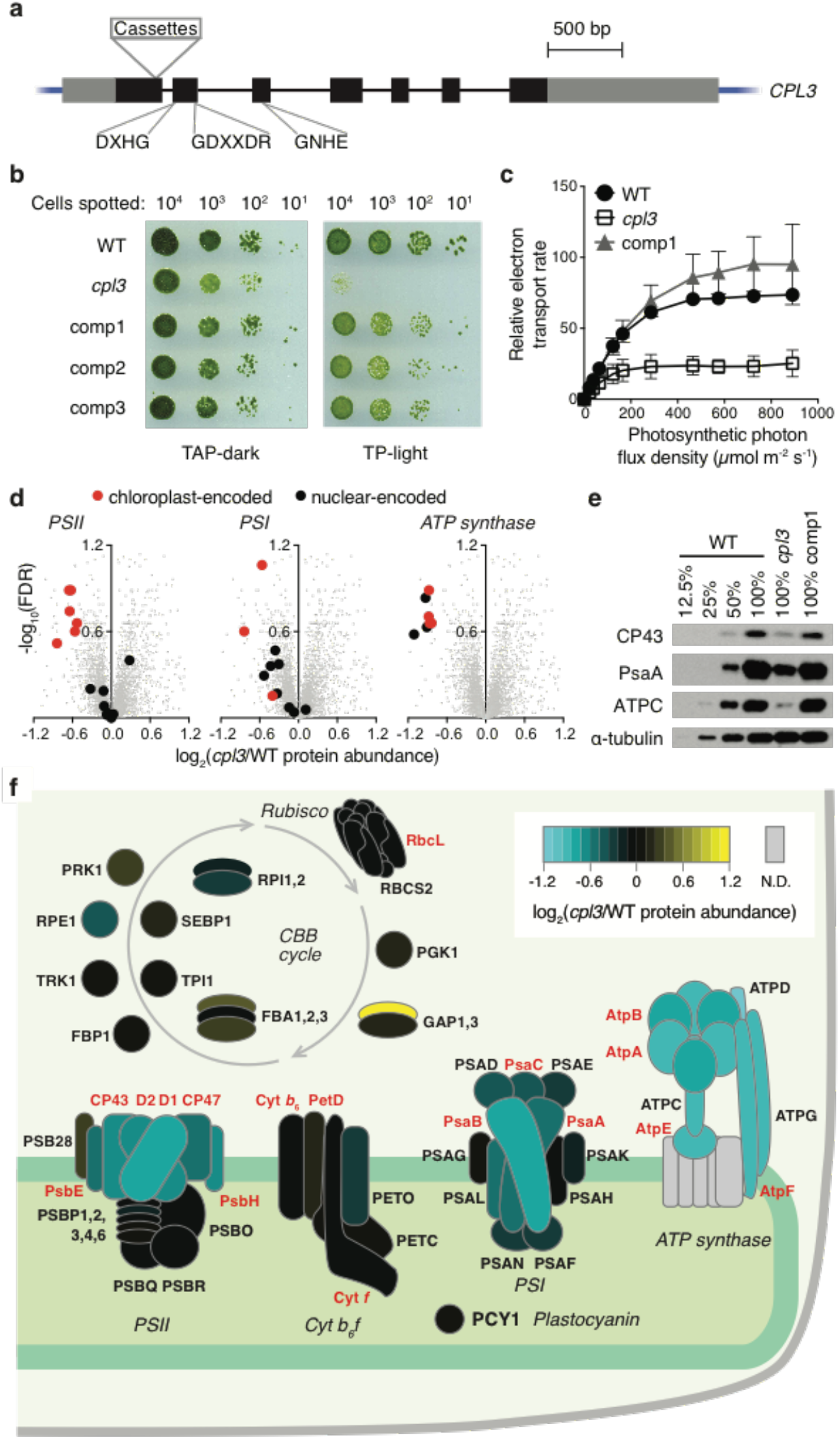
CPL3 is required for photosynthetic growth and accumulation of photosynthetic protein complexes in the thylakoid membranes. **a**, The *cpl3* mutant contains cassettes inserted in the first exon of *CPL3*. The locations of conserved protein phosphatase motifs are indicated (see Supplementary Fig. 6e). Black boxes indicate exons; gray boxes indicate UTRs. **b**, *cpl3* is deficient in growth under photosynthetic conditions. The growth deficiency is rescued upon complementation with a wild-type copy of the *CPL3* gene (comp1-3 represent three independent complemented lines). **c**, *cpl3* has a lower relative photosynthetic electron transport rate than the wild-type strain (WT) and comp1. Error bars indicate standard deviation (n ≥ 3). **d**, Whole-cell proteomics indicate that *cpl3* is deficient in the accumulation of PSII, PSI, and the chloroplast ATP synthase. Each gray dot represents one Chlamydomonas protein. The subunits of PSII, PSI and ATP synthase are highlighted as black or red symbols. See also Supplementary Table 14. **e**, Western blots show that CPL3 is required for normal accumulation of the PSII subunit CP43, the PSI subunit PsaA, and the chloroplast ATP synthase subunit ATPC. α-tubulin was used as a loading control. **f**, A schematic summary of the protein abundance of subunits in the light reactions protein complexes or enzymes in the CBB cycle in *cpl3* relative to the wild type based on proteomics data. The relative abundance is shown as a heatmap. Depicted subunits that were not detected by proteomics are filled with gray. Nuclear-encoded proteins are labeled in black font while chloroplast-encoded subunits are labeled in red font. A stack of horizontal ovals indicates different isoforms for the same enzyme, such as FBA1, FBA2, and FBA3. Cyt, cytochrome.

Consistent with the pooled growth data, *cpl3* showed a severe defect in photosynthetic growth on agar, which was rescued under heterotrophic conditions (Fig. 4b). We confirmed that the *CPL3* gene is disrupted in the *cpl3* mutant and found that complementation with a wild-type copy of the *CPL3* gene rescues the phenotype, demonstrating that the mutation in *CPL3* is the cause of the growth defect of the mutant (Supplementary Note and Supplementary Fig. 6a-d).

We then examined the photosynthetic performance, morphology of the chloroplast, and the composition of photosynthetic pigments and proteins in *cpl3*. Photosynthetic electron transport rate was decreased under all light intensities, suggesting a defect in the photosynthetic machinery (Fig. 4c). The chloroplast morphology of *cpl3* appeared similar to the wild type based on chlorophyll fluorescence microscopy (Supplementary Fig. 7a). However, we observed a lower chlorophyll *a*/*b* ratio in *cpl3* than in the wild type (Supplementary Fig. 7b), which suggests a defect in the accumulation or composition of the protein-pigment complexes involved in the light reactions^21^. Using whole-cell proteomics, we found that *cpl3* was deficient in accumulation of all detectable subunits of the chloroplast ATP synthase (ATPC, ATPD, ATPG, AtpA, AtpB, AtpE, AtpF), some subunits of PSII (D1, D2, CP43, CP47, PsbE, PsbH), and some subunits of PSI (PsaA and PsaB) (FDR<0.31 for each subunit, Fig. 4d, Fig. 4f, and Supplementary Table 14). We confirmed these findings by western blots on CP43, PsaA, and ATPC (Fig. 4e). Our results indicate that CPL3 is required for normal accumulation of thylakoid protein complexes (PSII, PSI, and ATP synthase) involved in the light reactions of photosynthesis.

Our finding that 21/43 of the higher-confidence photosynthesis hit genes were uncharacterized suggests that nearly half of the genes required for photosynthesis remain to be characterized. This finding is remarkable, considering that genetic studies on photosynthesis extend back to the 1950s^22^. Our validation of *CPL3*’s role in photosynthesis illustrates the value of the uncharacterized hit genes identified in this study as a rich set of candidates for the community to pursue.

More broadly, it is our hope that the mutant resource presented here will serve as a powerful complement to newly developed gene editing techniques^23-28^, and that together these tools will help the research community generate fundamental insights in a wide range of fields, from organelle biogenesis and function to organism-environment interactions.

## Acknowledgments

We thank Olivier Vallon for helpful discussions; Matthew Cahn and Garret Huntress for developing and improving the CLiP website; Xuhuai Ji at the Stanford Functional Genomics Facility and Ziming Weng at the Stanford Center for Genomics and Personalized Medicine for deep sequencing services; Alan Itakura for help in library pooling; Shriya Ghosh, Kyssia Mendoza, and Matthew LaVoie for technical assistance; Kathryn Barton, Winslow Briggs, and Zhi-Yong Wang for providing lab space; Joseph Ecker, Liz Freeman Rosenzweig and Moshe Kafri for constructive suggestions on the manuscript; and the Princeton Mass Spectrometry Facility for proteomics services. This project was supported by a grant from the National Science Foundation (MCB-1146621) awarded to M.C.J. and A.R.G., grants from the National Institutes of Health (DP2-GM-119137) and the Simons Foundation and HHMI (55108535) awarded to M.C.J., a German Academic Exchange Service (DAAD) research fellowship to F.F., Simons Foundation fellowships of the Life Sciences Research Foundation to R.E.J. and J.V.-B., EMBO long term fellowship (ALTF 1450-2014 and ALTF 563-2013) to J.V.-B and S.R., and a Swiss National Science Foundation Advanced PostDoc Mobility Fellowship (P2GEP3_148531) to S.R.

## Author contributions

X.L. developed the method for generating barcoded cassettes; R.Y. and S.R.B. optimized the mutant generation protocol; R.Y., N.I., and X.L. generated the library; J.M.R., N.I., A.G., and R.Y. maintained, consolidated, and cryopreserved the library; X.L. developed the barcode sequencing method; N.I., X.L., R.Y., and W.P. performed combinatorial pooling and super-pool barcode sequencing; X.L. performed LEAP-Seq; W.P. developed mutant mapping data analysis pipeline and performed data analyses of barcode sequencing and LEAP-Seq; W.P. analyzed insertion coverage and hot/cold spots; R.Z. and J.M.R. performed insertion verification PCRs and Southern blots; F.F., R.E.J., and J.V.-B. developed the library screening protocol; F.F., J.V.-B., and X.L. performed the photosynthesis mutant screen and barcode sequencing; R.E.J. and W.P. developed screen data analysis methods and implemented them for the photosynthesis screen; X.L. and T.M.W. annotated the hits from the photosynthesis screen; X.L., J.M.R., and S.R. performed growth analysis, molecular characterizations, and complementation of *cpl3*; S.S. and T.M.W. performed physiological characterizations of *cpl3*; M.T.M. and S.S. performed western blots on the photosynthetic protein complexes; M.T.M. performed microscopy on *cpl3*; X.L., W.P., and T.S. performed proteomic analyses; M.L. and P.A.L. maintained, cryopreserved, and distributed mutants at the Chlamydomonas Resource Center; X.L., W.P., A.R.G., and M.C.J. wrote the manuscript with input from all authors; M.C.J. and A.R.G. conceived and guided the research and obtained funding.

## Supplementary Information

Methods

Supplementary Note

Supplementary Figures 1-7

Supplementary Tables 1-14 (as separate excel or text files)

## SUPPLEMENTARY INFORMATION

### Other supplementary materials for this manuscript (provided as separate excel or text files)

Supplementary Table 1. Primers and experimental design for all PCRs related to library generation and mapping.

Supplementary Table 2. Binary codes for plate super-pooling.

Supplementary Table 3. Binary codes for colony super-pooling.

Supplementary Table 4. Read counts for each barcode in each combinatorial super-pool.

Supplementary Table 5. List of all mapped mutants in the library. Supplementary Table 6. Primers and results of PCRs used to verify the insertion sites of randomly-picked mutants from the mutant library.

Supplementary Table 7. Statistically significant insertion hot spots and cold spots. Supplementary Table 8. Statistically significant depleted functional terms. Supplementary Table 9. Candidate essential genes.

Supplementary Table 10. Read counts of barcodes before and after pooled growth in the photosynthesis screen.

Supplementary Table 11. Statistics of the pooled growth data for all genes. Supplementary Table 12. Summary of previous characterizations of higher- and lower-confidence genes’ roles in photosynthesis.

Supplementary Table 13. Read counts of *cpl3* exon and intron alleles in the pooled screens.

Supplementary Table 14. Proteomic characterization of the *cpl3* mutant.

### Methods

#### Generation of the indexed and barcoded mutant library: a conceptual overview

A three-step pipeline was developed for the generation of an indexed, barcoded library of insertional mutants in Chlamydomonas (Fig. 1b and Supplementary Fig. 1).

To generated mutants, CC-4533^1^ (“wild type” in text and figures) cells were transformed with DNA cassettes that randomly insert into the genome, confer paromomycin resistance for selection, and inactivate the genes they insert into. Each cassette contained two unique 22 nucleotide barcodes, one at each end of the cassette (Supplementary Fig. 1a-d). Transformants were arrayed on agar plates and each insertion in a transformant would contain two barcodes. The barcode sequences as well as the insertion site were initially unknown (Supplementary Fig. 1e).

To determine the sequences of the barcodes in each colony, combinatorial pools of the individual mutants were generated, with DNA extracted, and barcodes amplified and deep-sequenced. The combinatorial pooling patterns were designed so that each colony was included in a different combination of pools, allowing us to determine the barcode sequences associated with individual colonies based on which pools the sequences were found in (Supplementary Fig. 1f and Supplementary Fig. 2a-e; Methods). This procedure was similar in concept to the approach we used in our pilot study^2^, but it consumed significantly less time because we used a simple PCR amplifying only the barcodes instead of a multi-step flanking sequence extraction protocol (ChlaMmeSeq^1^) on each combinatorial pool.

To determine the insertion site associated with each barcode, the library was pooled into a single sample or six separate samples. The barcodes and their flanking genomic DNA were PCR amplified using LEAP-Seq^2^ (Supplementary Fig. 1g and Supplementary Fig. 2f-j; Methods). The flanking sequences associated with each barcode were obtained by paired-end deep sequencing^3,4^. The final product is an indexed library in which each colony has known flanking sequences that identify the genomic insertion site, and barcode sequences that facilitate pooled screens in which individual mutants can be tracked by deep sequencing (Fig. 3a).

Experimental details for this pipeline are described in paragraphs below.

#### Generation of insertion cassettes

The insertion cassette designated <u>C</u>assette containing <u>I</u>nternal <u>B</u>arcodes <u>1</u> (CIB1) was generated in four steps: (1) generating double-stranded DNAs containing random sequences (Supplementary Fig. 1a); (2) digesting the double-stranded DNAs to yield cassette ends (Supplementary Fig. 1a); (3) obtaining the backbone from digestion of plasmid pMJ016c that contains the sequences between the two barcodes (Supplementary Fig. 1b); (4) ligating the two cassette ends with the cassette backbone (Supplementary Fig. 1c).

Step 1: To generate each end of the cassette that contains barcodes, a long oligonucleotide primer (Supplementary Fig. 1a and Supplementary Table 1) containing a random sequence region of 22 nucleotides was used as a template for the extension of a shorter oligonucleotide primer. Each 50-µL reaction mixture contained 32 µL H_2_O, 10 µL Phusion GC buffer, 1.5 µL DMSO, 1 µL 10 mM dNTP, 2.5 µL 10 µM long oligo, 2.5 µL 10 µM short oligo, and 0.5 µL Phusion HS II DNA polymerase (F549L, Thermo Fisher). The reaction mixtures were subjected to a single thermal cycle: 98°C for 40 sec, 97°C to 63°C ramp (−1°C every 10 sec), 63°C for 30 sec, 72°C for 5 min.

Step 2: The double-stranded product yielded from Step 1 was digested using *Bsa*I (R0535L, New England Biolabs). For the 5’ side primer extension product, the digestion yielded two bands of 87 bp (plus 4 nt of overhang) and 31 bp (plus 4 nt of overhang). For the 3’ side, they were 68 bp and 31 bp. The larger band from each digestion was purified from a 2.5% agarose gel using D-tubes (71508-3, EMD Millipore) as previously described^1^ (Supplementary Fig. 1a).

Step 3: The synthesized plasmid pMJ016c, which contains the *HSP70-RBCS2* promoter, the paromomycin resistance gene *AphVIII*, and the *PSAD* and *RPL12* terminators, was digested using *Bsa*I. Two bands of 2064 bp and 3363 bp were obtained. The 2064 bp band (cassette backbone) was purified from a 0.8% agarose gel using the QIAquick Kit (28106, Qiagen) according to the manufacturer’s instructions (Supplementary Fig. 1b).

Step 4: The two fragments and the cassette backbone were ligated using T4 DNA ligase (M0202L, New England Biolabs) (Supplementary Fig. 1c). Each 30-µL reaction mixture contained 38 ng 5’ cassette end, 30 ng 3’ cassette end, 305 ng cassette backbone, 3 µL ligase buffer, and 0.5 µL ligase. The double-stranded product of 2,223 bp was gel purified using D-tubes and used for mutant generation. The sequence of the CIB1 cassette generated (Supplementary Fig. 1d) has been uploaded to the mutant ordering website: https://www.chlamylibrary.org/showCassette?cassette=CIB1.

#### Mutant generation, mutant maintenance, and medium recipes

Chlamydomonas CC-4533 strain was grown in Tris-Acetate-Phosphate (TAP) medium in a 20-L container under 100 μmol photons m^-2^ s^-1^ light (measured at the periphery) to a density of 1-1.5x 10^6^ cells/mL. Cells were collected by centrifugation at 300-1,000*g* for 4 min. Pellets were washed once with 25 mL TAP medium supplemented with 40 mM sucrose, and then resuspended in TAP supplemented with 40 mM sucrose at 2x 10^8^ cells/mL. 250 µL of cell suspension was then aliquoted into each electroporation cuvette (Bio-Rad) and incubated at 16°C for 5-30 min. For each cuvette, 5 μL DNA cassette CIB1 at 5 ng/μL was added to the cell suspension and mixed by pipetting. Electroporation was performed immediately as previously described^1^. After electroporation, cells from each cuvette were diluted into 8 mL TAP supplemented with 40 mM sucrose and shaken gently in dark for 6 h. After incubation, cells were plated on TAP containing 20 µg/mL paromomycin (800 µL per plate) and incubated in darkness for approximately two weeks before colony picking.

Approximately 210,000 total mutants were picked using a Norgren CP7200 colony picking robot and maintained on 570 agar plates, each containing a 384-colony array. We propagated this original, full library by robotically passaging the mutant arrays to fresh 1.5% agar solidified TAP medium containing 20 μg/mL paromomycin using a Singer RoToR robot (Singer Instruments)^2^. The full collection was grown in complete darkness at room temperature and passaged every four weeks. In this collection, 127,847 of the mutants were mapped. Colonies that failed to yield barcodes or flanking sequences may contain truncated insertion cassettes^1^ that have lost the primer binding sites used for barcode amplification or LEAP-Seq analysis. By removing the mutants that were not mapped, mutants that did not survive propagation, and some of the mutants in genes with 20 or more insertions, we consolidated 62,389 mutants into 245 plates of 384-colony arrays for long-term robotic propagation.

The TAP medium was prepared as previously reported^5^. The TP medium used in this research was similar to TAP except that HCl instead of acetic acid was used to adjust the pH to 7.5.

#### Combinatorial pooling

For combinatorial pooling and barcode determination for each mutant colony, 570 plate-pools (each containing all mutants on one plate) and 384 colony-pools (each containing all mutants in the same colony position across all plates) were generated from two separate sets of the library as previously described^2^. Binary error-correcting codes were used to design combinatorial pooling schemes, as previously described^2^. The existence of suitable binary error-correcting codes and their mathematical construction methods were checked using an online database^6^. For colony super-pooling, the same 384-codeword subset of the [20,10,6] code as previously employed^2^ was used. For plate super-pooling, the [21,11,6] code was generated by triple shortening of the [24,14,6] code^7^. In order to ensure detection of cases of two colonies derived from a single mutant, which could otherwise cause incorrect colony locations to be identified for such mutants, the subset of codewords with a bit sum of 10 (708 codewords) was taken from the [21,11,6] code, using the choose_codewords_by_bit_sum function. Both subsets of codewords were checked for the possibility of such sister colony conflicts using the clonality_conflict_check function: no conflicts were detected up to 2 errors, meaning any incorrect result due to a sister colony case would have at least 2 differences compared to any expected correct result. The final subset of 570 codewords for plate super-pooling was chosen as previously^2^. The final codeword lists are provided as Supplementary Tables 2 and 3.

Generation of plate-super-pools and colony-super-pools from the plate-pools and colony-pools was performed using the Biomek FX liquid handling robot (Beckman Coulter) as previously described^2^. The instruction files for the Biomek robot were generated using the robotic_plate_transfer.py program.

#### Barcode amplification from super-pools

DNA was extracted from super-pool samples as previously described^1^ and the barcodes were amplified (Supplementary Fig. 1f) using the Phusion HSII PCR system. For either 5’ or 3’ barcode amplifications, one primer (5’ R1 or 3’ R1; sequences provided in Supplementary Table 1) used in the PCR was common for all super-pools; the other primer (5’ R2-1, 5’ R2-2,…; 3’ R2-1, 3’ R2-2,…;) contained an index sequence that allows multiplexed sequencing, i.e. combining of multiple samples in one sequencing lane. Each 50 μL PCR mixture contained 125 ng genomic DNA, 10 μL GC buffer, 5 μL DMSO, 1 μL dNTPs at 10 mM, 1 μL (for 5’) or 2 μL (for 3’) MgCl_2_ at 50 mM, 2.5 μL of each primer at 10 μM, and 1 μL Phusion HSII polymerase. The reaction mixtures were incubated at 98°C for 3 min, followed by 10 three-step cycles (10 sec at 98°C, 25 sec at 58°C or 63°C for 5’ and 3’ barcodes respectively, and 15 sec at 72°C), and then 8 two-step cycles (10 sec at 98°C, and 40 sec at 72°C). Similar amount of products from three to eight super-pools were combined, purified using MinElute columns (28006, Qiagen), and the product bands (235 bp for 5’ and 209 bp for 3’) were gel purified. The purified products were sequenced using the Illumina HiSeq platform from a single end with a custom primer (5’ Seq and 3’ Seq, Supplementary Table 1).

#### Deconvolution of super-pool sequencing data

The barcode sequences were extracted from the Illumina sequencing data from each super-pool using the cutadapt command-line program^8^, with a 13 bp expected cassette sequence, allowing 1 alignment error, and taking the trimmed barcode reads between 21 and 23 bp in length. The command for 5’ sequences was “cutadapt -a GGCAAGCTAGAGA -e 0.1 -m 21 -M 23”, and for 3’ sequences “cutadapt -a TAGCGCGGGGCGT -e 0.1 -m 21 -M 23”. A barcode was found in 97-99% of the sequences in each super-pool.

The reads for each distinct barcode sequence in each super-pool were counted (Supplementary Table 4). Many of the sequenced barcodes are likely to contain PCR or sequencing errors. Such barcodes were left uncorrected, because they are very unlikely to appear in enough super-pools to be deconvolved and included in the final data. The deconvolution based on the read count table was performed as previously described^2^, for 5’ and 3’ data separately. A single set of optimized (N, x) parameters was chosen for each dataset, with m = 0 in all cases: N = 8 and x = 0.14 for 5’ plate-super-pool data, N = 8 and x = 0.16 for 3’ plate-super-pool data, N = 6 and x = 0.12 for 5’ colony-super-pool data, N = 6 and x = 0.1 for 3’ colony-super-pool data. Note that data for colony-super-pool 14 are missing for plates 351-570, which caused imperfections in the deconvolution process, but the missing data were dispensable due to the error-correction capability built into the pooling scheme.

#### LEAP-Seq

To connect the flanking sequence with the corresponding barcode for each insertion, we performed LEAP-Seq as reported before^2^ except that barcodes in addition to the flanking sequences were included in the amplicons (Supplementary Fig. 1g, and Supplementary Fig. 2f). Genomic DNA of mutants in the library was used as the template for the extension of a biotinylated primer that anneals to the insertion cassette. The primer extension products were purified by binding to streptavidin-coupled magnetic beads and then ligated to a single-stranded DNA adapter. The ligation products were then used as templates for PCR amplification. The PCR products were gel-purified before being submitted for deep sequencing.

We tried different combinations of primers and attempted to perform LEAP-Seq either on six sub-pools (each containing mutants from one-sixth of the library) separately or on the entire library in a single reaction (Supplementary Table 1). Sequencing results from all the samples were used in the analyses below.

#### Basic LEAP-Seq data analysis

The LEAP-Seq samples were sequenced with Illumina Hi-Seq, yielding paired-end reads. Each read pair has a proximal side, containing the barcode, a part of the cassette sequence, and the immediate genomic flanking sequence; and a distal side, containing the genomic sequence a variable distance away (Supplementary Fig. 2f-j).

A newly developed method was used to separate cassette sequence from the proximal reads and thus identify the barcode and genomic flanking sequence even in cases where the cassette was truncated. This was done using the deepseq_strip_cassette.py script, which uses local bowtie2 alignment^9^ to detect short cassette sequence. A bowtie2 alignment was performed against the expected cassette sequence (GGAGACGTGTTTCTGACGAGGGCTCGTGTGACTAGTGAGTCCAAC for 5’ reads and ACTGACGTCGAGCCTTCTGGCAGACTAGTTGCTCCTGAGTCCAAC for 3’ reads), using the following bowtie2 options: “–local –all –ma 3 –mp 5,5 –np 1 –rdg 5,3 –rfg 4,3 –score-min C,20,0 -N0 -L5 -i C,1,0 -R5 -D30 –norc –reorder”. The alignments for each proximal read were filtered to only consider cases where the cassette aligns after a 21-23 bp barcode, at most 5 bp of expected initial cassette sequence are missing, and at least 10 bp of expected cassette sequence are aligned with at most 30% errors. Out of the filtered alignments, the best one was chosen in a maximally deterministic manner, in order to ensure that multiple reads of the same insertion junction yield the same result. The alignment with the highest alignment score is chosen (the bowtie scoring function was customized to distinguish between as many cases as possible); if there were multiple alignments with the same score, the one with the longer alignment was chosen.

The resulting cassette alignment was then removed from each proximal read, with the section before the cassette being considered the barcode and the section after the cassette being considered the genomic flanking region. The resulting genomic proximal reads and the raw genomic distal reads were trimmed to 30 bp using the fastx_trimmer command-line utility (http://hannonlab.cshl.edu/fastx_toolkit), aligned to the Chlamydomonas genome (version 5.5 from Phytozome^10^) and the cassette, and the alignments were filtered to yield a single result using deepseq_alignment_wrapper.py, as previously described^1^.

The barcode sequences and proximal and distal alignment results were merged into a single dataset, with data grouped into insertion junctions based on the barcode, using the add_RISCC_alignment_files_to_data function. Data relating to barcodes that were not present in the combinatorial deconvolution results were discarded. The gene-related information for each insertion junction was added using the find_genes_for_mutants and add_gene_annotation functions. All functions in this paragraph are methods of the Insertional_mutant_pool_dataset class in the mutant_IB_RISCC_classes.py module.

#### Detecting pairs of flanking sequences that correspond to two sides of the same insertion (confidence levels 1 or 2)

Pairs of insertion junctions likely derived from two sides of the same insertion were identified using the deconvolution_utilities.get_matching_sides_from_table function, using the method previously described^2^, with an additional distance bin of 1-10 kb. The resulting pair counts were as follows:

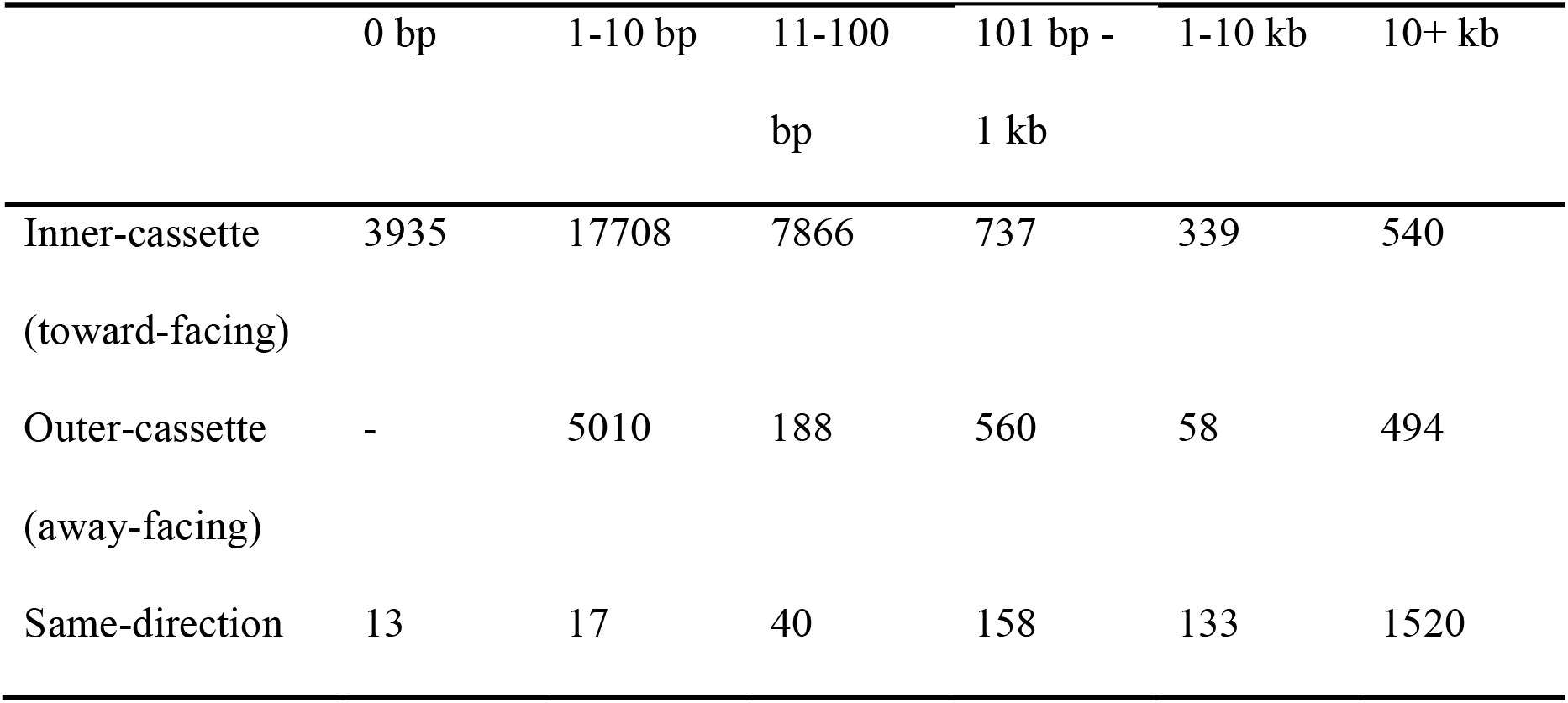

Additionally, there were 22,247 pairs in which the two junctions were mapped to different chromosomes.

The number of inner-cassette pairs is significantly larger than 50% of the number of same-direction pairs in all size ranges up to 10 kb, implying that most of the inner-cassette pairs in those size ranges are derived from a single insertion with a genomic deletion corresponding to the distance. This can be further confirmed by looking at the indicators of the probability of correct mapping for the insertion junctions: insertions with both sides mapped to the same region are almost certainly correctly mapped, and therefore independent indications of their correct mapping should be higher than for other insertions. As expected, the inner-cassette pairs up to 10 kb have a higher fraction of very high confidence insertion pairs (with both sides having 70% or more read pairs mapping to the same locus, and 500 bp or higher longest distance spanned by such read pairs): for size ranges up to 10 kb, 37-41% of the pairs are very high confidence, while for 10+ kb the number is only 16%.

The number of outer-cassette pairs is significantly larger than 50% of the number of same-direction pairs in size ranges between 1 bp and 1 kb, implying that most of the outer-cassette pairs in those size ranges are derived from a single insertion. There are two possible physical interpretations of a single insertion yielding an outer-cassette pair of insertion junctions: (1) an insertion with a genomic duplication causing the same genomic DNA sequence to be present on both sides of the cassette (potentially due to single-strand repair); and (2) an insertion of two cassettes flanking a “junk” fragment of genomic DNA. The 1-10 bp cases must be a genomic duplication, since a 1-10 bp “junk” fragment could not yield a 30 bp flanking sequences aligning to the genome. This is confirmed by 41% of the pairs being very high confidence. The 101 bp-1 kb cases are almost certainly insertions of two cassettes flanking a “junk” fragment, based on only 3.8% of them being very high confidence. The 188 11-100 bp cases, with a 27% very high confidence, are likely split between the two categories; based on previous analysis^1^ we used 30 bp as the cutoff between cases 1 and 2 for outer-cassette pairs. The case 2 pairs, i.e. insertions of two cassettes flanking a junk fragment, were used to determine the typical range of lengths of junk fragments (Supplementary Fig. 3f).

Based on this analysis, all insertion junction pairs likely to be derived from two sides of the same insertion (inner-cassette up to 10 kb and outer-cassette up to 30 bp) were categorized as confidence level 1 (extremely likely to be correctly mapped) because their mapping position is derived from two independent flanking sequences. They were annotated in Supplementary Table 5 as confidence level 1, and the “if_both_sides” column was set to “perfect” for the 0 bp distance cases, “deletion” for the remaining inner-cassette cases, and “duplication” for the outer-cassette cases.

A similar type of analysis was performed to look for pairs of insertion junctions derived from two sides of an insertion with a junk fragment. For each pair of insertion junctions in one colony (except pairs of insertion junctions already identified as two sides of the same insertion), we looked at the distance and relative orientation between the proximal read of the first junction and each distal read from the second junction; cases where the distal read was mapped to within 10 kb of the proximal read were counted as matches. We repeated the process with the first and second junctions swapped. To simplify the analysis, two cases were ignored: colonies with matches between more than two insertions (~12% of match cases), and insertion pairs where the proximal read of one insertion was a match to multiple distal reads of the other insertion with different orientations (~3% of match cases). We then took the distance to the closest distal read, and counted the cases by orientation and distance, as before:

|  | 0-10 bp | 11-100 bp | 101 bp - 1<br>kb | 1-10 kb |
| --- | --- | --- | --- | --- |
| Inner-cassette (toward-facing) | 11 | 5072 | 5787 | 289 |
| Outer-cassette (away-facing) | 28 | 140 | 152 | 82 |
| Same-direction | 6 | 185 | 283 | 195 |

Note that the distances are expected to be higher in this case, because if we are looking at a case of two sides of one insertion with a junk fragment, the distal read will be a variable distance away from the junk-genome junction which is the actual insertion location. So even for insertions with no genomic deletion/duplication, the distance between the proximal read on one side and the nearest distal read on the other side will not be 0 bp.

The number of inner-cassette cases up to 1 kb is more than 10x larger than the number of same-direction cases, so these insertion pairs are extremely likely to be two sides of one insertion with a junk fragment (and possibly a genomic deletion). Thus, all the pairs in this category were identified as confidence level 2, which are extremely likely to be correctly mapped.

The number of inner-cassette cases with a distance of 1-10 kb and the number of outer-cassette cases with a distance of 0-10 bp is also higher than the expected 50% of the same-direction cases, suggesting that many of them are also two sides of the same insertion, but the differences are less dramatic and thus the number of false positives would be too high for us to be comfortable identifying all these pairs as confidence level 2.

The insertion position information for junk fragment sides of confidence level 2 insertions originally reflected the junk fragment rather than the actual genomic insertion position. We corrected it to show the nearest distal read matching the non-junk side: the flanking sequence and position was changed to that of that distal read; the “LEAPseq_distance” field was changed to the longest distance between two distal reads that mapped to the presumed real insertion position (i.e. to the same region as the proximal read of the insertion junction from the other side); the remaining LEAPseq fields were likewise changed to reflect the numbers of distal reads and positions mapped to the presumed real insertion position. For confidence level 2 insertions, the “if_both_sides” column was set to “with-junk”; for the sides with a junk fragment, the “if_fixed_position” column was set to “yes_nearest_distal”, and for the sides without a junk fragment it was kept as “no”.

The confidence level 1 and 2 insertions (counting only the non-junk side of the confidence level 2 insertions) appear to be of high quality (Supplementary Fig. 2h).

#### Categorizing the remaining insertions and correcting junk fragments (confidence levels 3 and 4)

After identifying the highest-confidence insertion junctions, i.e. those with two matching sides of the same insertion, we sought to separate the remaining insertions (with only one side mapped) into a set with a high likelihood of having correctly mapped genomic insertion positions and a set with insertion positions likely to reflect junk fragments. We considered two factors to separate these two sets: (1) the percentage of read pairs that map to the same locus, and (2) the longest distance spanned by such a read pair (Supplementary Fig. 2, i and j). We decided to solely use the first factor based on the fact that nearly all of the insertions with low distances but high percentage of read pairs mapped to the same locus were ones with relatively few LEAP-Seq reads, indicating that their short distances spanned are likely due to them having few reads (and thus a lower chance of a long read) rather than to a junk fragment. Therefore we decided to use the percentage of read pairs mapping to the same locus as the only factor in distinguishing the higher and lower confidence insertion sets, because that factor is independent of the number of reads. To determine what cutoff would be appropriate, we took advantage of the already known confidence level 1 insertions. We calculated the fraction of confidence level 1 pairs among all the colonies with exactly two insertions (two insertions are required for a confidence level 1 pair) as an approximate lower bound on the number of correctly mapped insertions. Over the entire dataset, this fraction is 65%; when calculated only on insertions with at least 50% read pairs mapping to the same locus, it’s 78%; for insertions with at least 60%, 70%, 80% and 90% read pairs mapping to the same locus, it is 79%. Thus it is clear that using a cutoff anywhere in the 50-90% range significantly improves the quality of the dataset, regardless of the exact position of the cutoff. This makes sense, because the 50-90% range constitutes a very small fraction of all insertions. We opted to use 60% as the cutoff for confidence level 3, i.e. insertions with only one mapped side but with LEAP-Seq data indicating very likely correct mapping.

The remaining insertions, with below 60% read pairs mapping to the same locus and thus with the proximal LEAP-Seq read likely to be part of a junk fragment, were analyzed further to identify the most likely true insertion position. The same analysis was applied to all insertions with the proximal LEAP-Seq read with no genomic alignment (possibly due to a very short junk fragment resulting in the 30 bp proximal read being a hybrid of the junk fragment sequence and genomic sequence from the real insertion position, or simply due to PCR or sequencing errors yielding an unmappable sequence), or with multiple equally good genomic alignments (which could be derived from the real genomic location, but in a non-unique region of the genome, requiring the use of distal reads to determine the correct insertion location), or mapped to the insertion cassette (indicating a second cassette fragment inserted between the first cassette and the genome, which can be treated the same way as a junk genomic DNA fragment).

In order to determine the best method of identifying the true insertion location based on the full distal LEAP-Seq read data, we grouped the distal LEAP-Seq reads for each insertion into regions no more than 3 kb in size. For each such group, we calculated three measures that we thought might be the best method of identifying the real insertion location: (1) the number of reads in the group, (2) the number of unique genomic positions to which reads in the group were mapped, and (3) the distance spanned by the reads. LEAP-Seq reads mapped to the insertion cassette, or with no unique mapping to the genome, were excluded. In order to determine which method was the best, we used the junk fragment sides of confidence level 2 insertions, since for those the distal reads corresponding to the true genomic insertion locations had already been determined by an independent method (i.e. by matching the proximal read of the other side of the insertion). For each of the three methods listed above, the insertion location predicted by the method was compared to the known insertion location of each confidence level 2 insertion with a junk fragment. The results were as follows: 90% of the known insertion positions were correctly predicted by taking the region with the most total distal reads, 84% by taking the region with the most unique mapping positions, and 84% by taking the region with the longest distance spanned by the reads. Thus, the total number of distal reads was chosen as the most likely measure to yield the correct genomic insertion position of insertions with a junk fragment.

This method was then applied to all the insertions listed in the previous paragraph, yielding the most likely true location for each insertion; insertions with only a single LEAP-Seq distal read in each region were excluded, because one read did not provide enough data to determine the insertion position with any confidence. For some insertions, the region with the most distal LEAP-Seq reads also included the proximal LEAP-Seq read - in those cases, the original insertion position based on the proximal LEAP-Seq read was left unchanged. It is still possible that this position reflects a relatively long junk fragment rather than the true genomic insertion position, but we did not have enough data to distinguish those cases from high confidence. Likewise, it is possible that the corrected position with the most distal LEAP-Seq reads that do not match the proximal read reflects a second long junk fragment inserted after the first junk fragment which contains the proximal read (we know that insertions with multiple junk fragments can happen), but given the limited length of Illumina-sequenced LEAP-Seq reads, we cannot detect those cases with certainty, and have to limit ourselves to finding putative insertion positions that have a reasonably high probability of being correct.

Additionally, it turned out that many corrected positions for insertions originally mapped to the insertion cassette did not appear to be high-quality, with only a small fraction of distal reads mapped to the putative real insertion position. After looking at several such cases in detail, we concluded that they had not been analyzed correctly. They had single LEAP-Seq reads mapped to multiple distant locations on many chromosomes, compared to 100+ reads mapped to many cassette locations, with the putative real insertion position identified due to two or three single LEAP-Seq reads mapped close together on one chromosome. The uniformly low read numbers of genome-mapped reads compared with the high read numbers of cassette-mapped reads led us to conclude that the genome-mapped reads were results of PCR or sequencing errors or other artifacts, rather than being derived from real LEAP-Seq products, which should usually yield more than one read. Thus, those appeared to be cases where no LEAP-Seq products sequenced past the additional cassette fragment - this could be expected, because the full cassette is >2.2 kb in length, whereas vanishingly few LEAP-Seq reads are over 1.5 kb. In contrast, junk genomic DNA fragments are mostly smaller than 500 bp and all identified ones were below 1 kb, so this problem would not be expected to be common in genomic junk fragment cases. Indeed a cluster of low-matching-read-percent insertions was not observed in the corrected insertion positions in that category. We decided to exclude this category of incorrectly mapped insertions by only including corrected originally cassette-mapped insertions if >50% of the distal LEAP-Seq reads mapped to the putative correct insertion location.

All the insertions included in the final results of this analysis were annotated as confidence level 4. The final confidence level 4 insertions are of a relatively high quality (Supplementary Fig. 2j). The positions, flanking sequences and LEAP-Seq data of the corrected confidence level 4 insertions in Supplementary Table 5 were changed to reflect the new insertion position, in the same way as for the junk fragment sides of the confidence level 2 insertions above. An additional complication of the new corrected insertion positions was presented by the fact that the position of the nearest distal LEAP-Seq read is always at some distance from the true insertion position, depending on the length of the LEAP-Seq read. We attempted to correct for this by using confidence level 1 insertions to determine the average distance between the proximal read (reflecting the true insertion position) and the nearest distal read, separately for 5’ and 3’ datasets, depending on the total number of LEAP-Seq reads for the insertion (binned into ranges: 1, 2, 3, 4-5, 6-10, 11-20, 21+ total reads). For each confidence level 4 insertion with a corrected position, the position was further adjusted by the average distance for the correct side and number of reads as calculated above. This distance was appended as a number to the value in the “if_fixed_position” field for each insertion in Supplementary Table 5.

#### Insertion verification PCR

The PCR reactions were performed in two steps to verify the insertion site^2^ (Supplementary Table 6): (1) Genomic locus amplification: genomic primers that are ~1 kb away from the flanking genomic sequence reported by LEAP-Seq were used to amplify the genomic locus around the flanking sequence. If wild type produced the expected PCR band but the mutant did not produce it or produced a much larger product, this indicated that the genomic locus reported by LEAP-Seq may be disrupted by the insertional cassette and we proceeded to the second step; (2) Genome-cassette junction amplification: one primer binding to the cassette (omj913, GCACCAATCATGTCAAGCCT, for the 5’ side and omj944, GACGTTACAGCACACCCTTG, for the 3’ side) and the other primer binding to flanking Chlamydomonas genomic DNA (one of the genomic primers from the first step) were used to amplify the genome-cassette junction. If the mutant produced a PCR band with expected size that was confirmed by sequencing but wild type did not produced the expected PCR band, we categorized this insertion as “confirmed.” In some mutants, genomic primers surrounding the site of insertion did not yield any PCR products in wild type or the mutant even after several trials, possibly due to incorrect reference genome sequence or local PCR amplification difficulties. These cases were grouped as “failed PCR” and were not further analyzed.

72 mutants (24 insertions each for confidence levels 1 and 2, confidence level 3 and confidence level 4) were chosen randomly from the library and tested. The genomic DNA template was prepared from a single colony of each mutant using the DNeasy Plant Mini Kit (69106, Qiagen). The PCRs were performed using the Taq PCR core kit (201225, Qiagen) as described before^1^. PCR products of the expected size were verified by Sanger sequencing.

#### Southern blotting

Southern blotting was performed as previously described in detail^2^. Genomic DNA was digested with *Stu*I enzyme (R0187L, New England Biolabs) and separated on a 0.7% Tris-borate-EDTA (TBE) agarose gel. The DNA in the gel was depurinated in 0.25 M HCl, denatured in a bath of 0.5 M NaOH, 1M NaCl, neutralized in a bath of 1.5 M Tris-HCl, pH 7.4, 1.5 M NaCl, and finally transferred onto a Zeta-probe membrane (1620159, Bio-Rad) overnight using the alkaline transfer protocol given in the manual accompanying the membrane. On the next day, the membrane was gently washed with saline-sodium citrate (2xSSC: 0.3 M NaCl, 0.03 M sodium citrate), dried with paper towel, and UV cross-linked twice using the Stratalinker1800 (Stratagene). For probe generation, the *AphVIII* gene on CIB1 was amplified using primers oMJ588 (GACGACGCCCTGAGAGCCCT) and oMJ589 (TTAAAAAAATTCGTCCAGCAGGCG). The PCR product was purified and labeled according to the protocol of Amersham Gene Images AlkPhos Direct Labeling and Detection System (RPN3690, GE Healthcare). The membrane was hybridized at 60°C overnight with10 ng probe/mL hybridization buffer. On the next day, the membrane was washed with primary and secondary wash buffers and then visualized using a CLXPosure film (34093, Thermo Fisher).

#### Analyses of insertion distribution and identification of hot/cold spots

A mappability metric was defined to quantify the fraction of all possible flanking sequences from any genomic region that can be uniquely mapped to that region^1^. Calculation of mappability, hot/cold spot analysis and simulations of random insertions were performed as described previously^1^, except that a 30 bp flanking sequence lengths instead of a mix of 20 bp and 21 bp was used (because we now use 30 bp flanking sequence data derived from LEAP-Seq, rather than 20/21 bp ChlaMmeSeq sequences), and the v5.5 Chlamydomonas genome instead of the v5.3 genome was used^10^. This analysis was done on the original full set of mapped insertions, to avoid introducing bias from the choice of mutants into the consolidated set. The hot/cold spot analysis was performed on confidence level 1 insertions only, to avoid introducing bias caused by junk fragments and their imperfect correction. The full list of statistically significant hot/cold spots is provided in Supplementary Table 7.

#### Identification of underrepresented gene ontology (GO) terms

For each GO category, we calculated the total number of insertions in all genes annotated with the GO term and the total mappable (mappability defined in the Supplementary Note) length of all such genes, and compared them to the total number of insertions in and total mappable length of the set of flagellar proteome genes^11^. We compared these numbers using Fisher’s exact test, and did correction for multiple comparisons^12^ to obtain the false discovery rate (FDR). This analysis was done on the original full set of mapped insertions to avoid introducing bias from the choice of mutants into the consolidated set. We decided to use the flagellar proteome as the comparison set because flagellar genes are very unlikely to be essential; we did not use intergenic insertions or the entire genome because we know that the overall insertion density differs between genes and intergenic regions. The statistically significant results are listed in Supplementary Table 8.

#### Prediction of essential genes

To predict essential genes in Chlamydomonas, we sought to generate a list of genes that have fewer insertions than would be expected randomly was generated. Among them, those with 0 insertion are considered candidate essential genes.

To achieve these, for each gene, we calculated the total number of insertions in that gene and the total mappable length of that gene, and compared them to the total number of insertions in and total mappable length of the set of flagellar proteome genes^11^, as what we have performed on each GO category. The resulting list of genes with statistically significantly fewer insertions than expected is discussed in the Supplementary Note and shown in Supplementary Table 9: this includes 203 genes with no insertions, and 558 genes with at least one insertion. However, only genes 5 kb or longer yield an false discovery rate (FDR) of 0.05 or less when they have no insertions-our overall density of insertions is not high enough to detect smaller essential genes.

#### Pooled Screens

Library plates that were replicated once every four weeks onto fresh medium were switched to a 2-week replication interval to support uniform colony growth before pooling. Cells were pooled from 5-days-old library plates: first, for each set of eight agar plates, cells were scraped using the blunt side of a razor blade (55411-050, VWR) and resuspended in 40 mL liquid TAP medium in 50-mL conical tubes. Second, cells clumps were broken up by pipetting, using a P200 pipette tip attached to a 10-mL serological pipette. In addition, cells were pipetted through a 100 µm cell strainer (431752, Corning). Third, these sub-pools were combined as the master pool representing the full library.

The master pool was washed with TP, and resuspended in TP. Multiple aliquots of 2 x 10^8^ cells were pelleted by centrifugation (1,000*g*, 5 min, room temperature) and the supernatant was removed by decanting. Some aliquots were used for inoculation of pooled cultures, whereas other aliquots were frozen at −80 °C as initial pool samples for later barcode extraction to enable analysis of reproducibility between technical replicates. For pooled growth, 20 L TAP or TP in transparent Carboy containers (2251-0050, Nalgene) were inoculated with the initial pool to a final concentration of 2x 10^4^ cells/mL. Cultures were grown under 22°C, mixed using a conventional magnetic stir bar and aerated with air filtered using a 1 µm bacterial air venting filter (4308, Pall Laboratory). The TAP culture was grown in dark. For the two replicate TP cultures, the light intensity measured at the surface of the growth container was initially 100 µmol photons m^-2^ s^-1^, and then increased to 500 µmol photons m^-2^ s^-1^ after the culture reached ~2x 10^5^ cells/mL. When the culture reached the final cell density of 2x 10^6^ cells/mL after 7 doublings, 2x 10^8^ cells were pelleted by centrifugation (1,000*g*, 5 min, room temperature) for DNA extraction and barcode sequencing.

#### Barcode sequencing and data analysis for pooled screens

Barcodes were amplified and sequenced using the Illumina HiSeq platform as performed on the combinatorial super-pools in library mapping (Supplementary Fig. 1f). Initial reads were trimmed using cutadapt version 1.7.1^8^. Sequences were trimmed using the command “cutadapt -a <seq> -e 0.1 -m 21 -M 23 input_file.gz -o output_file.fastq”, where seq is GGCAAGCTAGAGA for 5’ data and TAGCGCGGGGCGT for 3’ data. Barcodes were counted by collapsing identical sequences using “fastx_collapser” (http://hannonlab.cshl.edu/fastx_toolkit). The barcode read counts for each dataset were normalized to a total of 100 million (Supplementary Table 10).

For evaluation of the quantitativeness of our barcode sequencing method, barcodes obtained from two technical replicate aliquots of the same initial pool were compared in read counts (Supplementary Fig. 5a). Barcodes obtained from the two TP-light cultures at the end of growth were compared to assess consistency between biological replicates (Fig. 3b).

To detect deficiency in photosynthetic growth, we compared mutant abundances in TP-light with TAP-dark at the end of growth (Fig. 3c). As a quality control, different barcodes in the same mutant were compared in the ratio of the TP-light read count to TAP-dark read count. Highly consistent ratios were observed (Supplementary Fig. 5b).

For the identification of photosynthetically deficient mutants, each barcode with at least 50 normalized reads in the TAP-dark dataset was classified as a hit if its ratio of normalized TP-light:TAP-dark read counts was 0.1 or lower, or a non-hit otherwise. The fraction of hit barcodes was 3.3% in replicate 1 and 2.9% in replicate 2. These barcodes represent 2,638 and 2,369 mutants showing a growth defect in the TP-light-I and TP-light-II replicates, respectively. A total of 3,109 mutants covering 2,599 genes showed a growth defect in either of the TP-light sample.

#### Identification and annotation of the hit genes from the screen

To evaluate the likelihood that a gene is truly required for photosynthesis, we counted the number of alleles for this gene with and without a phenotype, including exon/intron/5’UTR insertions. If the insertion was on the edge of one of those features, or in one of the features in only one of the splice variants, it was still counted. We excluded alleles with insertions in the 3’ UTRs, which we observed to less frequently cause a phenotype (Fig. 3, d and e). In cases of multiple barcodes in the same mutant (likely two sides of one insertion), the one with a higher TAP-dark read count was used for the calculation of normalized TP-light:TAP-dark read counts, to avoid double-counting a single allele. For each gene, a *P* value was generated using Fisher’s exact test comparing the numbers of alleles in that gene with and without a phenotype to the numbers of all insertions in the screen with and without a phenotype (Supplementary Table 11). A false discovery rate (FDR) correction was performed on the *P* values using the Benjamini-Hochberg method^12^, including only genes with at least 2 alleles present in the screen. Thus, genes with a single allele have a *P* value but lack a FDR.

This process was performed for both TP-light replicates. The list of higher-confidence genes was generated by taking genes with FDR of 0.27 or less in either replicate - this threshold includes all genes with 2 hit alleles and 0 non-hit alleles. The resulting list of hits included 37 genes in replicate 1, 34 in replicate 2, 44 total. The FDR values for the higher-confidence genes in both replicates are shown in Tables 1 and 2. Additionally, the list of lower-confidence genes was generated by taking genes with a *P* value of 0.058 or less – this value was chosen to include genes with only one allele with a phenotype and no alleles without a phenotype, but to exclude genes with one allele with and one without a phenotype. The resulting list included 264 genes total (210 in replicate 1, 196 in replicate 2).

**Table 1.**
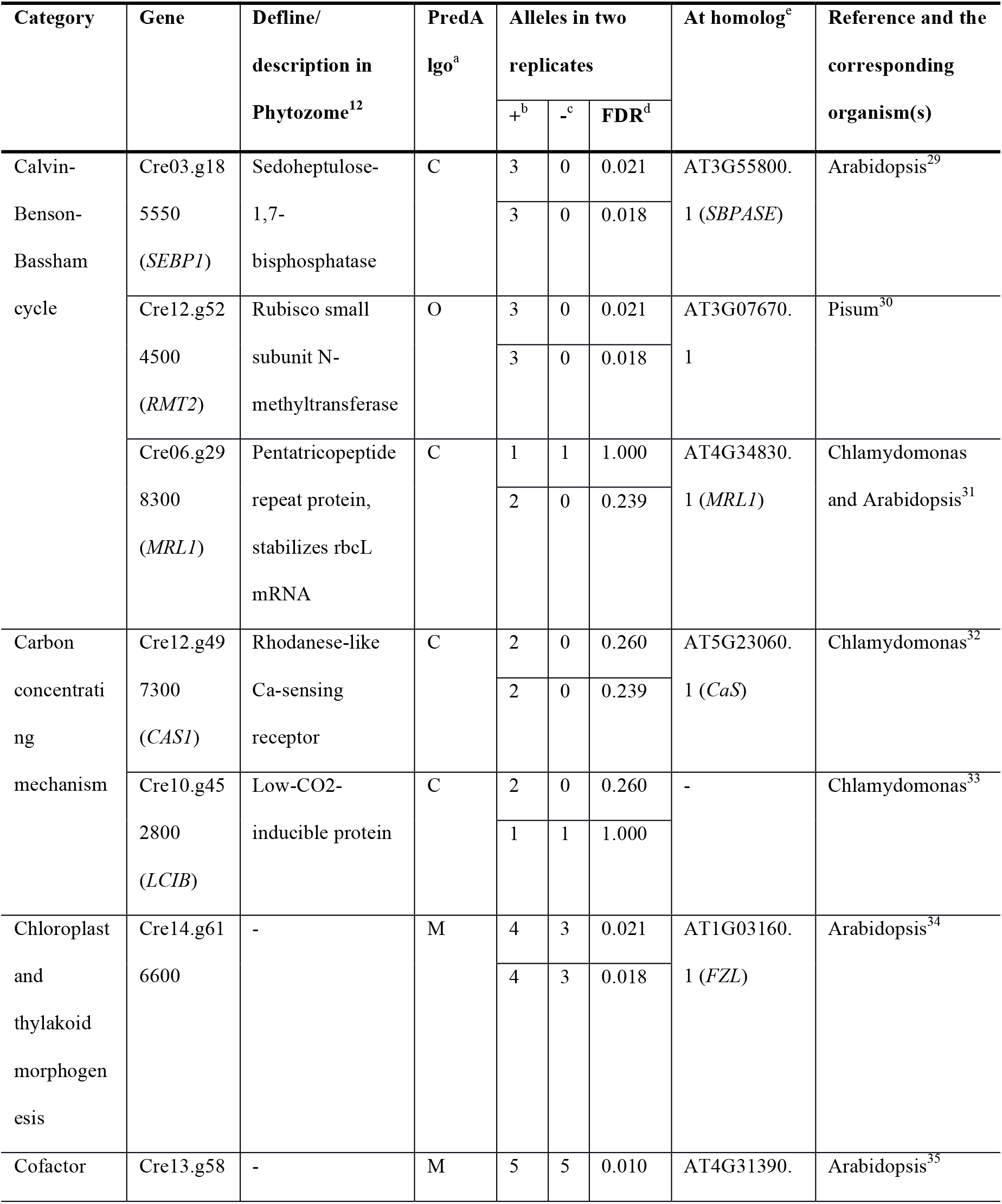
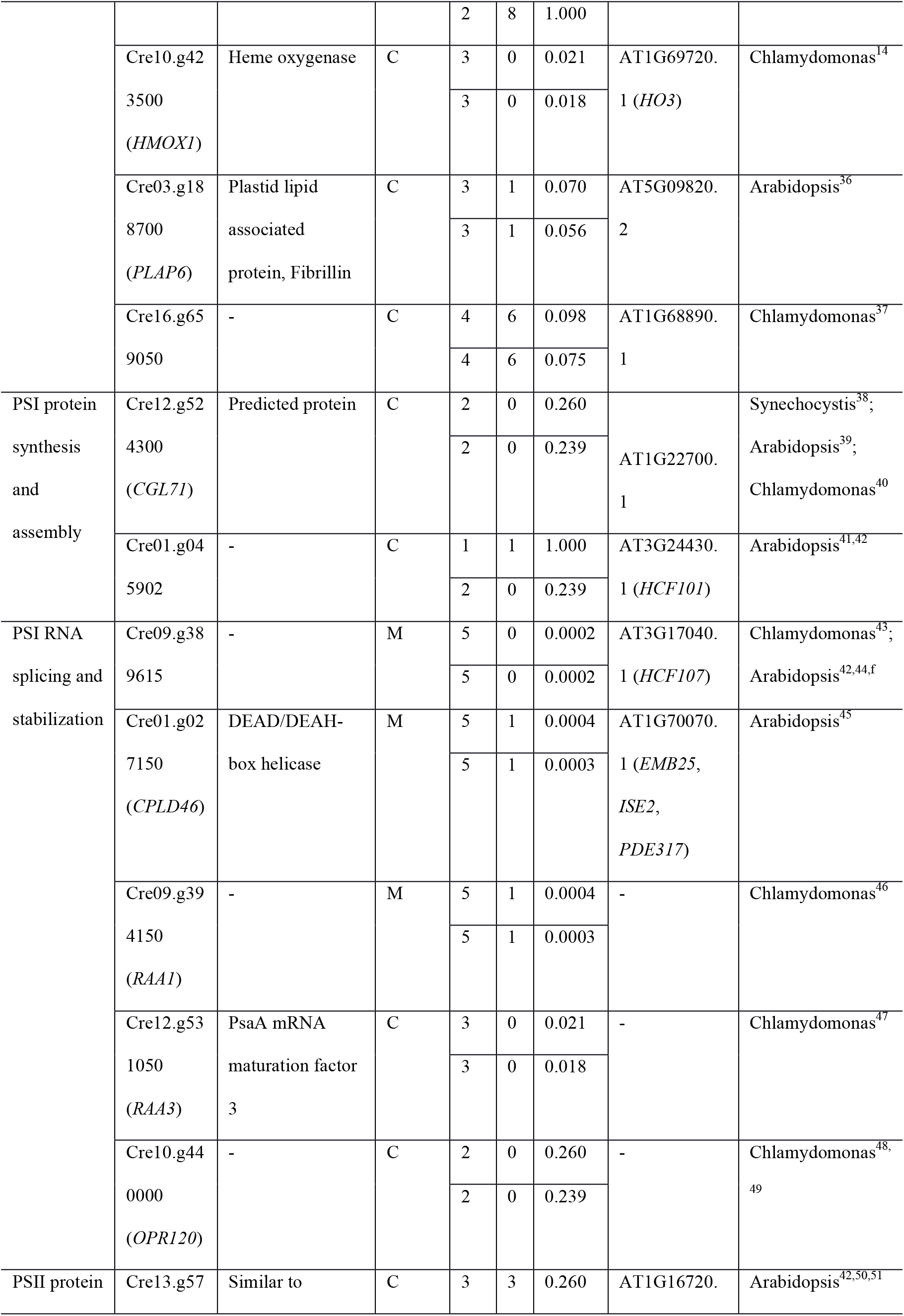
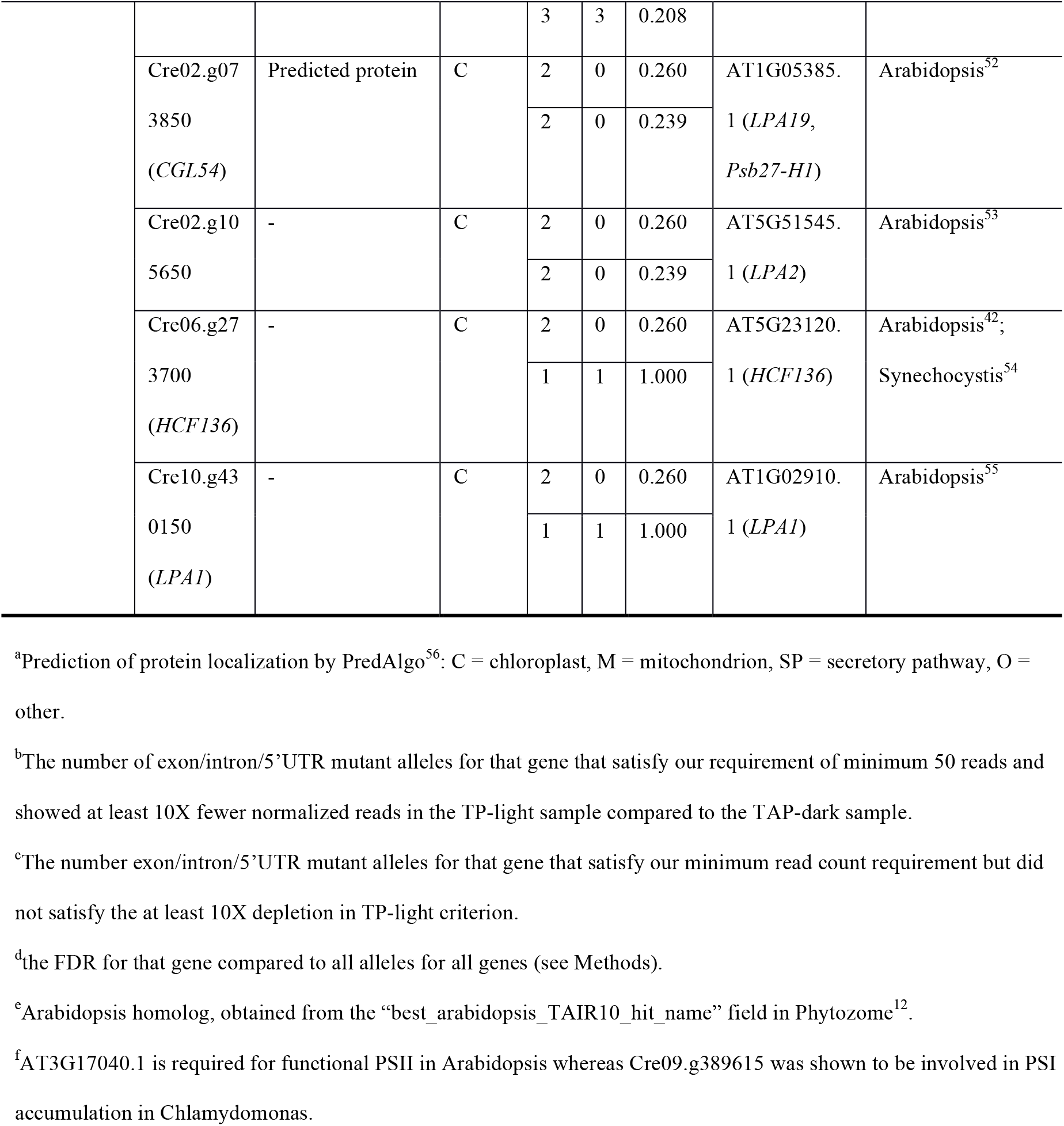
Higher-confidence genes from the photosynthesis screen that had a previously known role in photosynthesis.

**Table 2.** Higher-confidence genes from the photosynthesis screen with no previously known role in photosynthesis.

| Gene | Defline/description in<br>Phytozome | PredAlgo | Alleles in two<br>replicates |  |  | At homolog |
| --- | --- | --- | --- | --- | --- | --- |
|  |  |  | + | - | FDR |  |
| Cre01.g008550 | Serine/threonine kinase-related | O | 2 | 0 | 0.260 | AT1G73450.1 |
|  |  |  | 1 | 1 | 1.000 |  |
| Cre01.g014000 | - | C | 3 | 0 | 0.021 | - |
|  |  |  | 3 | 0 | 0.018 |  |
| Cre01.g037800<br>(*TRX21*) | ATP binding protein; thioredoxin<br>domain | O | 3 | 3 | 0.260 | AT2G18990.1<br>(*TXND9*) |
|  |  |  | 1 | 5 | 1.000 |  |
| Cre02.g073900 | All-trans-10'-apo-beta-carotenal<br>13,14-cleaving dioxygenase | C | 3 | 1 | 0.070 | AT4G32810.1<br>(*ATCCD8*, *CCD8*,<br>*MAX4*) |
|  |  |  | 3 | 1 | 0.056 |  |
| Cre02.g111550 | Serine/threonine kinase-related | SP | 10 | 8 | $< 10^{-6}$ | AT4G24480.1 |
|  |  |  | 6 | 12 | 0.015 |  |
| Cre03.g185200<br>(*CPL3*) | Metallophosphoesterase/metallo-<br>dependent phosphatase | C | 3 | 4 | 0.260 | AT1G07010.1 |
|  |  |  | 3 | 4 | 0.239 |  |
| Cre06.g259100 | - | C | 1 | 4 | 1.000 | - |
|  |  |  | 3 | 2 | 0.117 |  |
| Cre06.g281800 | Domain of unknown function<br>(DUF1995) | C | 3 | 0 | 0.021 | - |
|  |  |  | 3 | 0 | 0.018 |  |
| Cre07.g316050<br>(*CDJ2*) | Chloroplast DnaJ-like protein | M | 2 | 0 | 0.260 | AT5G59610.1 |
|  |  |  | 1 | 1 | 1.000 |  |
| Cre07.g341850<br>(*EIF2*) | Translation initiation factor IF-2,<br>chloroplastic | C | 2 | 0 | 0.260 | AT1G17220.1 (*FUG1*) |
|  |  |  | 2 | 0 | 0.239 |  |
| Cre08.g358350<br>(*TDA1*) | Fast leu-rich domain-containing<sup>a</sup> | C | 3 | 2 | 0.152 | - |
|  |  |  | 3 | 2 | 0.117 |  |
| Cre09.g396250 | Phosphatidate cytidyltransferase | SP | 2 | 0 | 0.260 | AT5G04490.1 (*VTE5*) |
|  |  |  | 1 | 1 | 1.000 |  |
| Cre10.g429650 | Alpha/beta hydrolase family<br>(Abhydrolase_5) | O | 2 | 0 | 0.260 | - |
|  |  |  | 1 | 1 | 1.000 |  |
| Cre10.g448950 | Nocturnin | C | 1 | 1 | 1.000 | AT3G58560.1 |
|  |  |  | 2 | 0 | 0.239 |  |
| Cre11.g467712 | Structural maintenance of<br>chromosomes smc family<br>member; starch-binding domain | M | 7 | 7 | 0.0003 | AT5G05180.1 |
|  |  |  | 7 | 7 | 0.0003 |  |
| Cre12.g542569 | Ionotropic glutamate receptor | O | 0 | 2 | 1.000 | AT1G05200.1<br>(*ATGLR3.4*, *GLR3.4*,<br>*GLUR3*) |
|  |  |  | 2 | 0 | 0.239 |  |
| Cre13.g566400<br>(*OPR55*) | Fast leu-rich domain-containing<sup>a</sup> | M | 4 | 2 | 0.018 | - |
|  |  |  | 4 | 2 | 0.015 |  |
| Cre13.g574000<br>(*GEF1*) | Voltage-gated chloride channel | O | 1 | 11 | 1.000 | AT5G26240.1<br>(*ATCLC-D*, *CLC-D*) |
|  |  |  | 4 | 8 | 0.144 |  |
| Cre13.g586750 | Transportin 3 and importin | O | 3 | 4 | 0.260 | AT5G62600.1 |
|  |  |  | 2 | 5 | 1.000 |  |
| Cre16.g658950 | - | C | 2 | 2 | 0.909 | - |
|  |  |  | 3 | 1 | 0.056 |  |
| Cre50.g761497 | Magnesium transporter mrs2<br>homolog, mitochondrial | M | 2 | 0 | 0.260 | AT5G22830.1<br>(*ATMGT10*, *GMN10*,<br>*MGT10*, *MRS2-11*) |
|  |  |  | 2 | 0 | 0.239 |  |

<sup>a</sup>The annotation of “fast leu-rich domain-containing” cannot be confirmed by BLASTp analysis at NCBI<sup>57</sup>.

One gene in the original higher-confidence list and four genes in the original lower-confidence list encode subunits of the plastidic pyruvate dehydrogenase. Mutants in these genes require acetate to grow because they cannot generate acetyl-CoA from pyruvate but can generate acetyl-CoA from acetate. This requirement for acetate, rather than a defect in photosynthesis, likely explains why mutants in this gene showed a growth defect in TP-light^13^. Removal of these genes led to a final list of 43 higher-confidence genes and 260 lower-confidence genes (Fig. 3f, Tables 1 and 2, and Supplementary Table 12).

We identified 65 (22 higher-confidence and 43 lower-confidence) out of the 303 hit genes as “known” genes based on genetic evidence: mutation of this gene in Chlamydomonas or another organism caused a defect in photosynthesis. Among the remaining 238 “candidate” genes (21 higher-confidence ones and 217 lower-confidence ones), some genes appear to be related to photosynthesis because of their predicted chloroplast localization or evolutionary conservation among photosynthetic organisms^14^, despite lack of solid genetic evidence. For three of the candidate genes (*CGL59*, *CPL3*, and *VTE5*), mutants with insertions adjacent to them were previously found to be acetate-requiring or hypersensitive to oxidative stress in the chloroplast^13^.

#### Analysis of candidate gene enrichment in reported transcriptional clusters related to photosynthesis

Two transcriptome datasets in Chlamydomonas were used in this analysis: a diurnal regulation study^15^ and a dark-to-light transition study^16^. For the first one, we chose the diurnal cluster 4 in the study that had photosynthesis-related genes enriched in it^15^. For the second one, we chose the genes upregulated upon transition to light^16^. In each case, the number of candidate genes included and not included in the regulated gene sets was compared to the total number of Chlamydomonas genes included and not included in the cluster, using Fisher’s exact test. The resulting *P* values were FDR-adjusted using the Benjamini-Hochberg method^12^.

#### Molecular characterization of the *cpl3* mutant

Mutant genotyping PCRs were performed as previously described^2^. Sequences of primers represented in Supplementary Fig. 6a are g1: CCGTCGTCACTTGC-TACAAC, g2: CGTAGTTGCAAGGGGTGTTT, c1: GACGTTACAGCACACCCTTG. To complement the *cpl3* mutant, the wild-type *CPL3* gene was PCR amplified and cloned into the vector pRAM118 vector that contains the aph7’’ gene^17^, which confers resistance to hygromycin B. In this construct, the expression of *CPL3* is under the control of the *PSAD* promoter. The construct was linearized before being transformed into the *cpl3* mutant. Transformants were robotically arrayed and assayed in colony sizes in the presence and absence of acetate respectively (Supplementary Fig. 6, c and d). Three representative lines that showed rescued photosynthetic growth were used in further phenotypic analyses (Fig. 4).

#### Analyses of growth, chlorophyll, and photosynthetic electron transport

For all physiological and biochemical characterizations of *cpl3* below, we grew cells heterotrophically in the dark to minimize secondary phenotypes due to defects in photosynthesis.

For spot assays, cells were grown in TAP medium in dark to log phase to around 10^6^ cells per mL. Cells were washed in TP and spotted onto solid TAP medium and TP medium respectively. The TAP plates were incubated in dark for 12 d before being imaged. The TP plates were incubated under 30 µmol photons m^-2^ s^-1^ light for 1 d, 100 µmol photons m^-2^ s^1^ light for 1 d, and then 500 µmol photons m^-2^ s^-1^ light for 4 d.

Chlorophyll *a* and *b* concentrations were measured as previously described^18^ using TAP-dark grown cells. We used TAP-dark-grown instead of TP-light-grown cells for chlorophyll analyses, photosynthetic performance analyses, microscopy, proteomics, and western blots (below) to avoid observing secondary effects due to the photosynthetic defects of the *cpl3* mutants.

To measure photosynthetic electron transport rate, TAP-dark grown cells were collected, re-suspended in fresh TAP medium, and dark acclimated for 20 min. Cells were then measured in chlorophyll fluorescence under a series of increasing light intensities using the “Light Curve” function on a DUAL-PAM-100 fluorometer (Walz). PSII quantum yield (ΦPSII) was quantified as previously described^19^. Relative electron transport rate (rETR) was calculated according to the following equation rETR = ΦPSII x I. I represents the emitted irradiance.

#### Microscopy

Cells were grown under the TAP-dark condition to log phase and concentrated ten-fold before microscopic analysis. Aliquots were deposited at the corner of a poly-L-lysine coated microslide well (Martinsried) and spread over the bottom of the well by overlaying with TAP-1% agarose at low temperature (<30°C), to minimize cell motion during image acquisition. Cells were imaged at room temperature though a Leica TCS SP5 laser scanning confocal microscope and an inverted 100x NA 1.46 oil objective. Chlorophyll fluorescence signal was generated using 514 nm excitation, and 650-690 nm collection. All images were captured using identical laser and magnification settings (4x zoom and single-slice through the median plane of the cell). Composite images (chlorophyll fluorescence overlay with bright field) were generated with Fiji^20^.

#### Proteomics

TAP-dark-grown cells were collected by centrifugation and flash-frozen. Proteins were extracted from the frozen pellets by resuspension in lysis buffer (6M guandium Hydrochloride, 10mM tris(2-carboxyethyl)phosphine, 40mM chloroacetamide, 100mM Tris pH8.5, 1x MS-Safe protease inhibitor, 1x Phosphatase inhibitor cocktail II), grinding with liquid nitrogen, followed by sonication. Protein lysates were then digested with trypsin (Promega) into peptides. Three biological replicates were processed for each strain.

The samples were labeled with tandem mass tags (TMTs), multiplexed and then fractionated before tandem mass spectrometry analyses. Briefly, each sample was labeled with the TMT labeling reagent (Thermo Fisher) according to the manufacturer’s instructions. The samples were then mixed in equimolar amounts and desalted using C18-stage tips^21^. The dried peptide mix was then separated using strong cation exchange (SCX) stage-tips^22^ into four fractions. Each of the four fractions were then diluted with 1% trifluoroacetic acid (TFA) and separated into three fractions using SDB-RPS stage tips. This procedure initially resulted in a total of 12 fractions. Fractions 1-3 (the children of the first SCX fraction) were pooled together yielding 10 final fractions. Each final fraction was diluted and injected per run using an Easy-nLC 1200 UPLC system (Thermo Fisher). Samples were loaded onto a nano capillary column packed with 1.9 µm C18-AQ (Dr. Maisch) mated to metal emitter in-line with a Fusion Lumos (Thermo Fisher). Samples were eluted using a split gradient of 10-20% solution B (80% ACN with 0.1% FA) in 32 min and 20-40% solution B in 92 min followed column wash at 100% solution B for 10 min. The mass spectrometer was operated in a data-dependent mode with the 60,000 resolution MS1 scan (380-1500 m/z), AGC target of 4e5 and max injection time of 50ms. Peptides above threshold 5e3 and charges 2-7 were selected for fragmentation with dynamic exclusion after 1 time for 60 s and 10 ppm tolerance. MS1 isolation windows of 1.6m/z, MS2 isolation windows 2 and HCD NCE of 55% were selected. MS3 fragments were detected in the Orbitrap at 50,000 resolution in the mass range of 120-500 with AGC 5e4 and max injection time of 86 ms. The total duty cycle was set to 3.0 sec.

Raw files were searched with MaxQuant^23^, using default settings for MS3 reporter TMT 10-plex data. Files were searched against sequences of nuclear, mitochondrial, and chloroplast-encoded Chlamydomonas proteins supplemented with common contaminants^10,24,25^. Raw files were also analyzed within the Proteome Discoverer (Thermo Fisher) using the Byonic^26^ search node (Protein Metrics). Data from Maxquant and Proteome Discoverer were combined in Scaffold Q+ (Proteome Software Inc.), which was used to validate MS/MS based peptide and protein identifications. Peptide identifications were accepted if they could be established at greater than 80.0% probability by the Scaffold Local FDR algorithm. Protein identifications were accepted if they could be established at greater than 96.0% probability and contained at least 2 identified peptides. Scaffold Q+ un-normalized data were exported in the format of the log_2_ value of the reporter ion intensities, which reflect the relative abundances of the same protein among different samples multiplexed. Each sample was then normalized to a median of 0 (by subtracting the original median from the raw values, since the values are log_2_). For each gene, for each pair of samples, the normalized log_2_ intensity values from the three replicates of one sample were compared against those for the other sample using a standard *t*-test. The resulting *P* values were adjusted for multiple testing^12^, yielding a false discovery rate (FDR) for each gene in each pair of samples. We note that our calculation of FDR does not take into account the spectral count of each protein (provided in Supplementary Table 14), which is related to the absolute abundance of the protein and impacts the accuracy of proteomic measurements. Specifically, proteins with a low spectral count are likely of low abundance in cells and often exhibit a large variation in the intensity value between the biological replicates.

#### Western blotting

TAP-dark grown cells were pelleted by centrifugation, resuspended in an extraction buffer containing 5 mM HEPES-KOH, pH 7.5, 100 mM dithiothreitol, 100 mM Na_2_CO_3_, 2% (w/v) SDS, and 12% (w/v) sucrose, and lysed by boiling for 1 min. Extracted proteins were separated on SDS-PAGE (12% precast polyacrylamide gels, BioRad) using tubulin as a loading and normalization control. Polypeptides were transferred onto polyvinylidene difluoride membranes using a semidry blotting apparatus (Bio-Rad) at 15 volts for 30 minutes. For western blot analyses, membranes were blocked for 1 h at room temperature in Tris-buffered saline-0.1% (v/v) Tween containing 5% powdered milk followed by a 1 h incubation of the membranes at room temperature with the primary antibodies in Tris-buffered saline-0.1% (v/v) Tween containing powdered milk (3% [w/v]). Primary antibodies were diluted according to the manufacturer’s recommendations. All antibodies were from Agrisera and the catalog numbers for the antibodies against CP43, PsaA, ATPC, and α-tubulin were AS11-1787, AS06-172-100, AS08-312, and AS10-680, respectively. Proteins were detected by enhanced chemiluminescence (K-12045-D20, Advansta) and imaged on a medical film processor (Konica) as previously described^2^.

#### Code availability

All programs written for this work are deposited at https://github.com/Jonikas-Lab/Li-Patena-2018.

### Supplementary Note

#### Accuracy of insertion mapping and number of insertions per mutant

In Chlamydomonas insertional mutants, short “junk fragments” of genomic DNA (likely from lysed cells) are often inserted between the cassette and flanking genomic DNA^1^. The difficulty in distinguishing these junk fragments from true flanking genomic DNA can lead to inaccurate mapping of the insertion to a genomic location^1,2^. Additionally, some cassettes are truncated during insertion, preventing mapping of the flanking sequence on one side. We sought to help users prioritize mutants for characterization by classifying insertions into categories that reflect our confidence in the mapping accuracy, based on two criteria: (1) whether flanking sequences from both sides of the cassette mapped to the same genomic region; and (2) whether the LEAP-Seq reads contained sequences from multiple genomic regions, suggesting the presence of junk DNA fragments inserted next to the cassette (Supplementary Fig. 3a and Supplementary Fig. 2f-j).

A confidence level of 1 was assigned to 19,015 insertions in which both cassette-genome junctions mapped to the same genomic region and were free of junk fragments. A confidence level of 2 was assigned to 5,665 insertions in which both cassette-genome junctions mapped to the same genomic region, after correcting for the presence of a junk fragment at one junction. A mapping confidence level of 3 was assigned to 36,600 insertions in which only one cassette-genome junction could be identified, with the likelihood of junk DNA insertion determined to be low based on fewer than 40% of LEAP-Seq reads containing sequence from multiple genomic regions. A mapping confidence level of 4 was assigned to 13,643 insertions in which only one junction could be identified, and that junction was likely to contain a junk fragment, or the flanking sequence could not be mapped to a unique genomic location. The mapping for these insertions was adjusted to reflect the most likely correct insertion site.

Approximately 95% of confidence level 1 and 2 insertions are mapped correctly based on PCR validation of randomly chosen mutants, compared to ~73% of confidence level 3 and ~58% of confidence level 4 (Supplementary Table 6; Methods).

Our bioinformatic analyses suggest that over 80% of the mutants harbor only one mapped insertion (Supplementary Fig. 3b), consistent with Southern blot data from randomly chosen mutants (Supplementary Fig. 3c).

#### Deletions, duplications, and junk fragments associated with insertions are small

Random insertions in Chlamydomonas are sometimes also associated with deletions and duplications of neighboring genomic DNA^13^. To further help users understand the quality of mutants in this library, we characterized these deletions and duplications by examining the sequences across both junctions of confidence level 1 insertions (Methods). Of these insertions, 11% had no deletions or duplications, 74% harbored genomic deletions and 15% had genomic duplications. The great majority (98%) of genomic deletions were less than 100 bp, but some were as large as 10 kb. While 98% of the genomic duplications were shorter than 10 bp, some extended to 30bp (Supplementary Fig. 3, d and e). Both the deletions and duplications likely resulted from non-homologous end joining repair that occurs during cassette insertion^27^. Additionally, examining the 651 insertions in which a junk fragment separated two cassettes inserted in the same location allowed us to estimate the typical junk fragment length. Most (73%) junk fragments were shorter than 300 bp, but some were as large as 1,000 bp (Supplementary Fig. 3f). If larger deletions, duplications or junk fragments were present, they were not sufficiently frequent to allow us to identify them reliably.

#### Insertion sites are randomly distributed with mild cold spots and a small number of hot spots

While a random insertion model produced a distribution of insertion sites broadly similar to the observed distribution (Fig. 1c and Supplementary Fig. 4a), we did detect some cold spots and hot spots where insertion density differed significantly from the random insertion model (Supplementary Fig. 4a; Supplementary Table 7; Methods). Cold spots cover 26% of the genome and on average show a 48% depletion of insertions. Hot spots cover 1.5% of the genome and contain 16% of insertions (Methods).

Hot spots fell into two distinct classes that differed in the local distribution of insertions (Supplementary Fig. 4, b and c). In one class, dozens of insertions were found within a region of 20-40 bp. In the other class, the insertions were distributed over a much larger region of 200-1,000 bp. Our observations suggest that hot spots could be caused by two distinct mechanisms; however, we did not observe a correlation between specific features of the genome (e.g. sequence, exon, intron, UTR, mappability) and the occurrence of either class of hot spots.

#### Absence of insertions identifies over 200 genes potentially essential for growth under the propagation conditions used

Identification of essential genes in bacteria, fungi, and mammals has revealed important molecular processes in these organisms^3,28-30^. We sought to take advantage of the very large set of mapped mutations in the library to identify candidate essential Chlamydomonas genes based on the absence of insertions in those genes (Methods). We note that our approach does not allow testing of gene essentiality under all possible conditions. Therefore, it is likely that some of the candidate essential genes we identify in this approach are required specifically for growth under our propagation conditions, but not under all conditions. For example, mutants in respiratory genes would be identified as essential if these mutants were not recovered under our propagation conditions (in the dark on acetate media), although the same mutants could have grown if recovery were under photosynthetic conditions.

Given our average density of insertions, we were able to detect a statistically significant (FDR< 0.05) lack of insertions for genes with a mappable length greater than 5 kb. We identified 203 candidate essential genes (Supplementary Table 9). We caution that this is a conservative list for two reasons: (1) if a gene has a mappable length smaller than 5 kb and has no insertion, its underrepresentation is not statistically significant; (2) some essential genes were not detected because there are insertions incorrectly mapped to them.

Many of these predicted essential genes have homologs that have been shown to be essential in other organisms. For example, Cre01.g029200 encodes a homolog of the yeast cell cycle protease separase ESP1^31^, Cre12.g521200 encodes a homolog of yeast DNA replication factor C complex subunit 1 RFC1^32^, and Cre09.g400553 encodes a homolog of the yeast nutrient status sensing kinase Target of Rapamycin 2 TOR2^33^. In addition, we observed genes encoding proteins involved in acetate utilization or respiration, such as acetyl-CoA synthetase/ligase^34^ (Cre07.g353450) and components of the mitochondrial F1F0 ATP synthase^35^ (Cre15.g635850 and Cre07.g340350). As discussed above, these genes may be essential under the conditions of library propagation, in which acetate serves as the energy source.

We also observed genes on the list with nonessential homologs in other organisms. One example is Cre13.g585301, which encodes monogalactosyldiacylglycerol (MGDG) synthase and whose Arabidopsis homolog MGD1 is not essential^36^. This can be explained by the presence of two other isoforms of MGDG synthases in Arabidopsis but not in Chlamydomonas^37^. Comparison of our candidate Chlamydomonas essential genes with those of other organisms can provide insights into evolutionary differences across the tree of life.

#### Deleterious mutations rather than differential chromatin configuration are the major cause of insertion density variation

One caveat for our above prediction of essential genes is that the lack of insertions could be caused by low chromatin accessibility at those loci to insertional mutagenesis. We reasoned that if chromatin accessibility influenced insertion density, the 3’ UTRs of these genes would also be less represented; while if low insertion density primarily reflected essentiality, we would still see many insertions in the 3’ UTRs of these genes, because 3’ UTR insertions typically do not disrupt gene function (Fig. 3, d and e). For all genes in the genome, we observed an insertion density of 1.1 insertions per mappable kb in exons and introns and 4.7 insertions per mappable kb in 3’ UTRs. For the candidate essential genes, despite a lack of insertions in exons and introns, the insertion density in 3’ UTRs is 4.1 insertions per mappable kb, similar to that of all genes. We thus conclude that low insertion density in our candidate essential genes is largely caused by mutations that impair mutant fitness instead of low chromatin accessibility to insertional mutagenesis.

#### Disruption of *CPL3* is the cause of the photosynthetic deficiency in the *cpl3* mutant

We sought to confirm and characterize the *cpl3* insertion in detail. Our high-throughput LEAP-Seq data suggested that *cpl3* contained an insertion of two back-to-back cassettes. Specifically, the *cpl3* mutant contains two insertion junctions from 3’ ends of two cassettes in opposite orientations, within the *CPL3* gene. Junction 1 is confidence level 3 (no junk fragment), and junction 2 is confidence level 4 (with a junk fragment, corrected) (Supplementary Fig. 6a). We successfully confirmed both junctions by PCR (Supplementary Fig. 6b). Sequencing of the product from junction 2 revealed that the end of the cassette has a 10-bp truncation and a 10-bp fragment of unknown origin inserted between the cassette and the *CPL3* gene. The genomic flanking sequence of junction 2 overlaps with the flanking sequence in junction 1 by 2 bp. When we amplified across the insertion site, *cpl3* yielded a product ~3 kb larger than the product from wild type (Supplementary Fig. 6b). Based on these results, the most likely model for this insertion is that two copies of the cassette (at least one truncated) inserted together into the *CPL3* gene in opposite orientations, with a 2-bp genomic duplication at the site of insertion.

To confirm the involvement of CPL3 in photosynthesis, we cloned *CPL3* genomic DNA and transformed it into the *cpl3* mutant. Based on colony size, photoautotrophic growth was rescued in approximately 14% of the transformants (Supplementary Fig. 6, c and d), a percentage consistent with previous Chlamydomonas genetic studies^38^. Three rescued transformants, named comp1-3, were chosen at random for phenotypic confirmation (Fig. 4b) and genotyping. In comp1-3, PCR across the insertion site of the *cpl3* mutation with primers “g1 + g2” yielded ~1.2 kb products (expected size: 1,311 bp) that indicate presence of wild-type *CPL3* sequence (from the wild-type *CPL3* in the complementation construct), and weak ~4 kb bands consistent with the presence of the original cassette insertion in *CPL3* (Supplementary Fig. 6b). The lower intensity of the ~4 kb bands in these samples can be explained by preferential amplification of the smaller template when multiple templates are present. To further confirm that comp1-3 still contained the original insertion in *CPL3*, we amplified the two insertion junctions in the complemented lines with primers “g1 + c1” and “g2 + c1”. These genetic complementation results demonstrate that the disruption of *CPL3* is the cause of the growth defect of the mutant.

### Supplementary Figures

**Supplementary Fig. 1.**
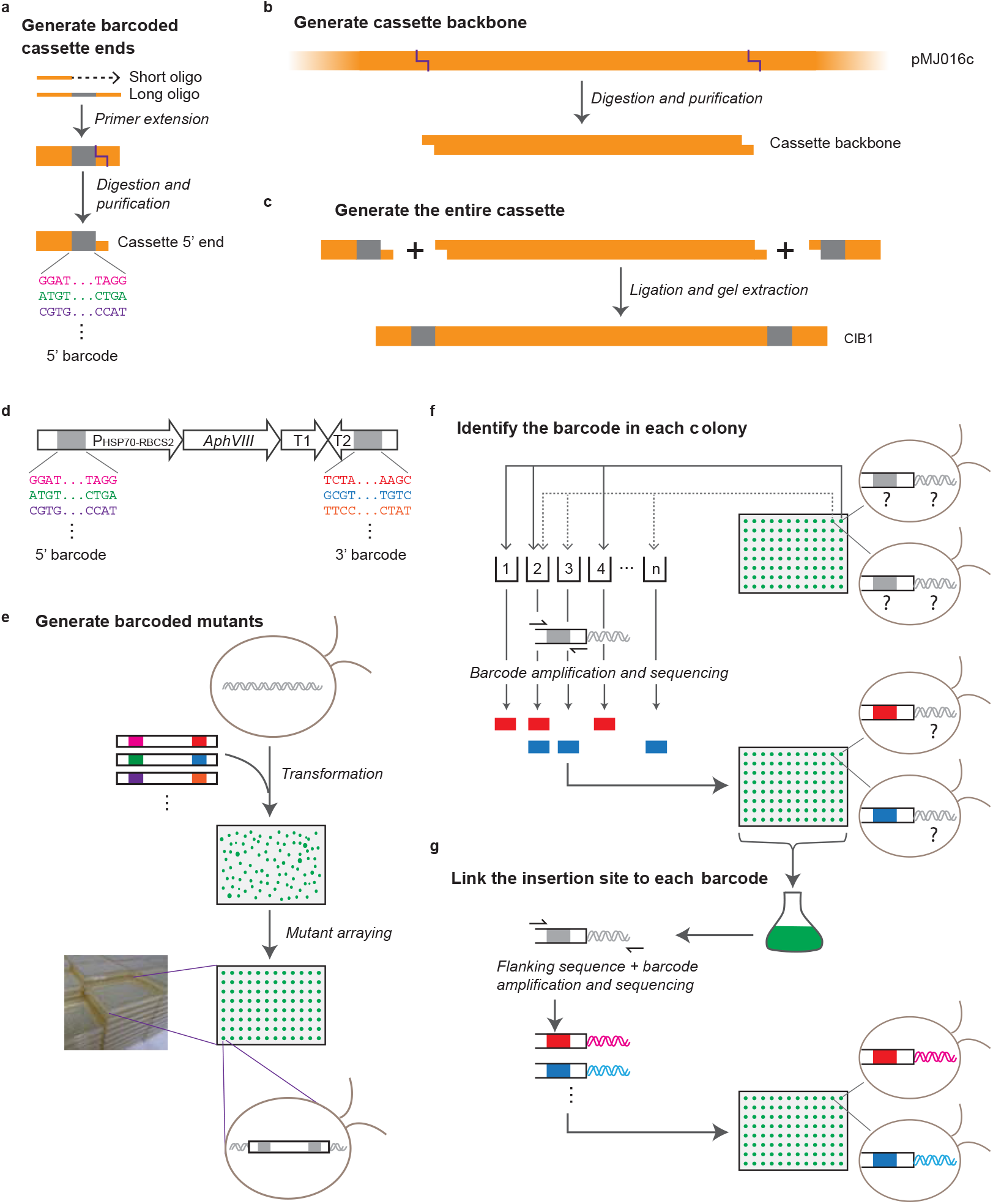
A pipeline was developed for generating barcoded cassettes (a-d) and for generating an indexed and barcoded library of insertion mutants in Chlamydomonas (e-g). **a**, A long oligonucleotide primer containing a random sequence region (indicated in gray) was used as a template for the extension of a shorter oligonucleotide primer (see Supplementary Table 1 for primer sequences). The resulting double-stranded product contains a random sequence region (22 bp in length; termed “barcode”). This product was restriction digested to generate a sticky end for subsequent ligation. The above steps were performed to produce both the 5’ and the 3’ ends of the cassette. The 5’ end of the cassette is shown as an example. **b**, The pMJ016c plasmid was digested to yield the backbone of the cassette. **c**, The 5’ and 3’ ends of the cassette generated above were ligated together with the cassette backbone to yield the cassette CIB1. **d**, The components of the cassette CIB1 are shown. CIB1 contains the *HSP70-RBCS2* promoter (with an intron from *RBCS2*), the *AphVIII* gene that confers resistance to paromomycin, two transcriptional terminators (T1: *PSAD* terminator; T2: *RPL12* terminator), and two barcodes (each 22 bp in length). **e**, Following transformation and arraying of individual mutants, the sequence of the barcodes contained in each insertion cassette was unique to each transformant but initially unknown for each colony. **f**, Barcodes were amplified from combinatorial pools of mutants, sequenced, and traced back to single colonies (Supplementary Fig. 2a-e; Methods). After this step, the barcode sequence for each colony was known. For simplicity, only one side of the cassette is shown. **g**, Barcodes and genomic sequences flanking the insertion cassettes were amplified from a pool of the library. By pooled next-generation sequencing, the sequence flanking each insertion cassette was paired with the corresponding barcode (Supplementary Fig. 2f). The flanking sequences were used to determine the insertion site in the genome. Because the colony location for each barcode was determined in the previous step, insertion sites could then be assigned to single colonies.

**Supplementary Fig. 2.**
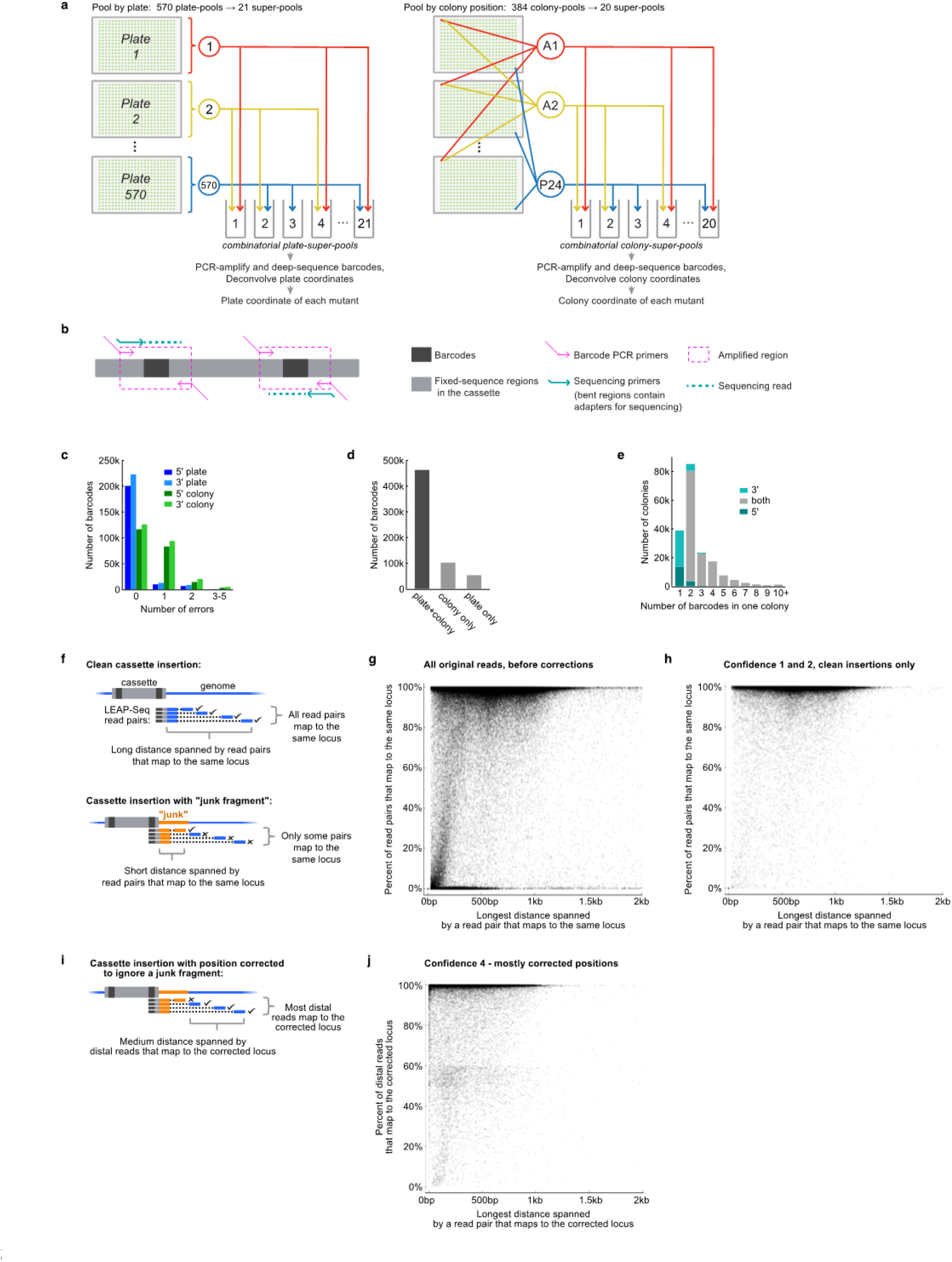
Combinatorial pooling, barcode deconvolution to colony, and determination of insertion sites. **a**, To determine which plate each barcode was on, each plate of mutants was pooled into one of 570 plate-pools. The plate-pools were then further combinatorially pooled into 21 plate-super-pools, in such a way that each plate-pool was in a unique combination of plate-super-pools. The barcodes present in each plate-super-pool were determined by deep sequencing, and the barcodes were assigned to plates based on the combination of plate-super-pools they were found in. A similar process was applied to the colony positions of each barcode. Combining the plate and colony data yielded a specific position for each barcode. **b**, The barcodes on the 5’ and 3’ sides of the cassette were sequenced separately, each with a single-end Illumina read. With the sequencing primers we used (indicated on the cassette), the reads start with the barcode sequence and extend into the cassette. **c**, Most barcode colony positions were identified with no errors, i.e. were found in one of the expected combinations of super-pools. Some were found in a combination of super-pools that had one or more differences from any expected combination, but the positions could still be identified due to the redundancy built into our method. The much higher number of one-error cases in the colony data compared to plate data is due to a loss of one of the colony-super-pools for a significant fraction of the samples (Methods). **d**, Both a plate and a colony position were identified for most barcodes. **e**, The number of barcodes mapped to an individual colony varied, with 2 being the most common. For colonies with two mapped barcodes, the large majority had one 5’ and one 3’ barcode, likely derived from two sides of one cassette. **f**, LEAP-Seq reads are paired-end reads with the proximal read containing the cassette barcode and immediate flanking genomic sequence, and the distal read containing flanking genomic sequence a variable distance away from the insertion site. During transformation, short fragments of genomic DNA, likely originating from lysed cells, are often inserted between the cassette and the true flanking genomic DNA. We refer to these short DNA fragments as “junk fragments”^1,2^. Such junk fragments can lead to incorrect insertion mapping if only the immediate flanking genomic sequence is obtained. LEAP-Seq data can be used to detect presence of junk fragments at an insertion junction based on two key characteristics: 1) the number of read pairs where both sides aligned to the same locus and 2) the longest distance spanned by such read pairs. **g**, The two key characteristics are plotted for the original full library, before any mapping corrections were applied. **h**, The same two characteristics are plotted for confidence level 1 and 2 insertions. For confidence level 2 insertions, only the side with no junk fragment is shown; for confidence level 1 insertions, one randomly chosen side is shown. **i**, LEAP-Seq data can be used to correct cases of probable junk fragment insertions and determine the most likely correct insertion position. The corrected data can be visualized using two modified key characteristics: the number of distal reads aligned to the corrected location, and the distance spanned by such reads. **j**, The modified characteristics are plotted for confidence level 4 insertions.

**Supplementary Fig. 3.**
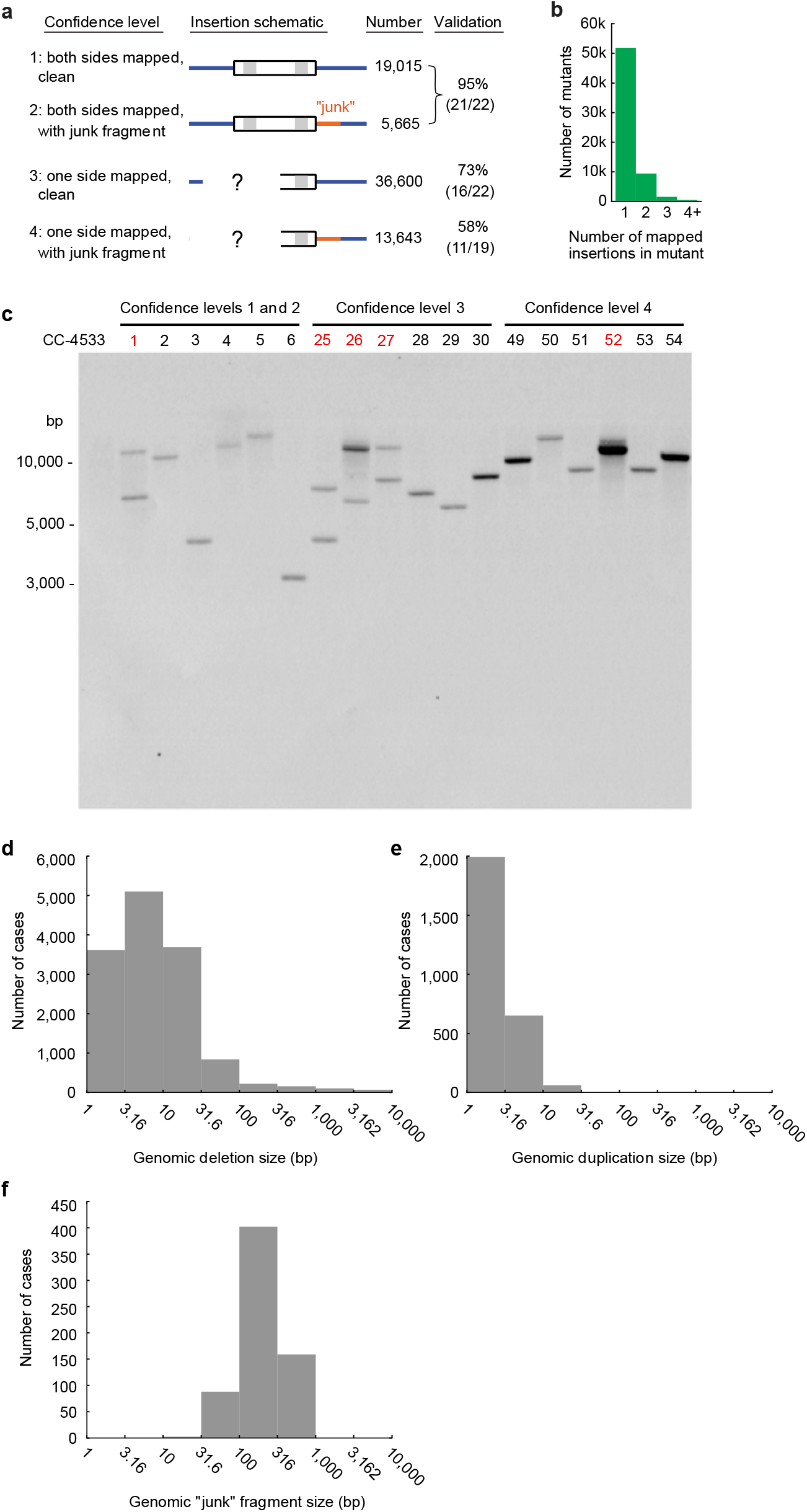
Characterization of genomic disruptions in mutants in the library. **a**, Mutants in the library were divided into four confidence levels, corresponding to different mapping scenarios. The insertion sites of a number of randomly chosen mutants in each category were verified by PCR (mutants from confidence levels 1 and 2 were assayed as one group; Supplementary Table 6). The numbers and percentages of confirmed insertions are shown in the last column. **b**, Most mutants have a single mapped insertion, and < 20% contain two or more mapped insertions. **c**, Eighteen randomly selected mutants from the four confidence levels were analyzed by Southern blotting using the coding sequence of *AphVIII* as the probe. Mutants are numbered and the details of their insertion sites are presented in Supplementary Table 6. The mutant number is highlighted in red when the Southern blot was interpreted to indicate at least two insertions in that mutant. The wild-type strain CC-4533 (WT) was included as a negative control. **d**, Most genomic deletions accompanying cassette insertions are smaller than 100 bp, but deletions up to 10 kb are present in some mutants. Deletions larger than 10 kb may also be present, but there were not enough of them to be clearly detected based on the aggregate numbers. **e**, Most genomic duplications accompanying cassette insertion are smaller than 10 bp, but they can be up to 30 bp. Larger duplications may be present, but these are not common enough to be detected based on the aggregate numbers. **f**, The distribution of junk fragment lengths was determined using a dataset of 651 insertions of two cassettes surrounding a junk fragment, allowing us to precisely map both ends of the junk fragment using LEAP-Seq. Most junk DNA fragments are smaller than 320 bp, but we have detected some up to 1 kb in size. Larger junk fragments may be present, but are not common enough to be detected based on the aggregate numbers. Note that the x-axes for **d-f** are set to the logarithmic scale. Data presented in this figure are described in the Supplementary Note.

**Supplementary Fig. 4.**
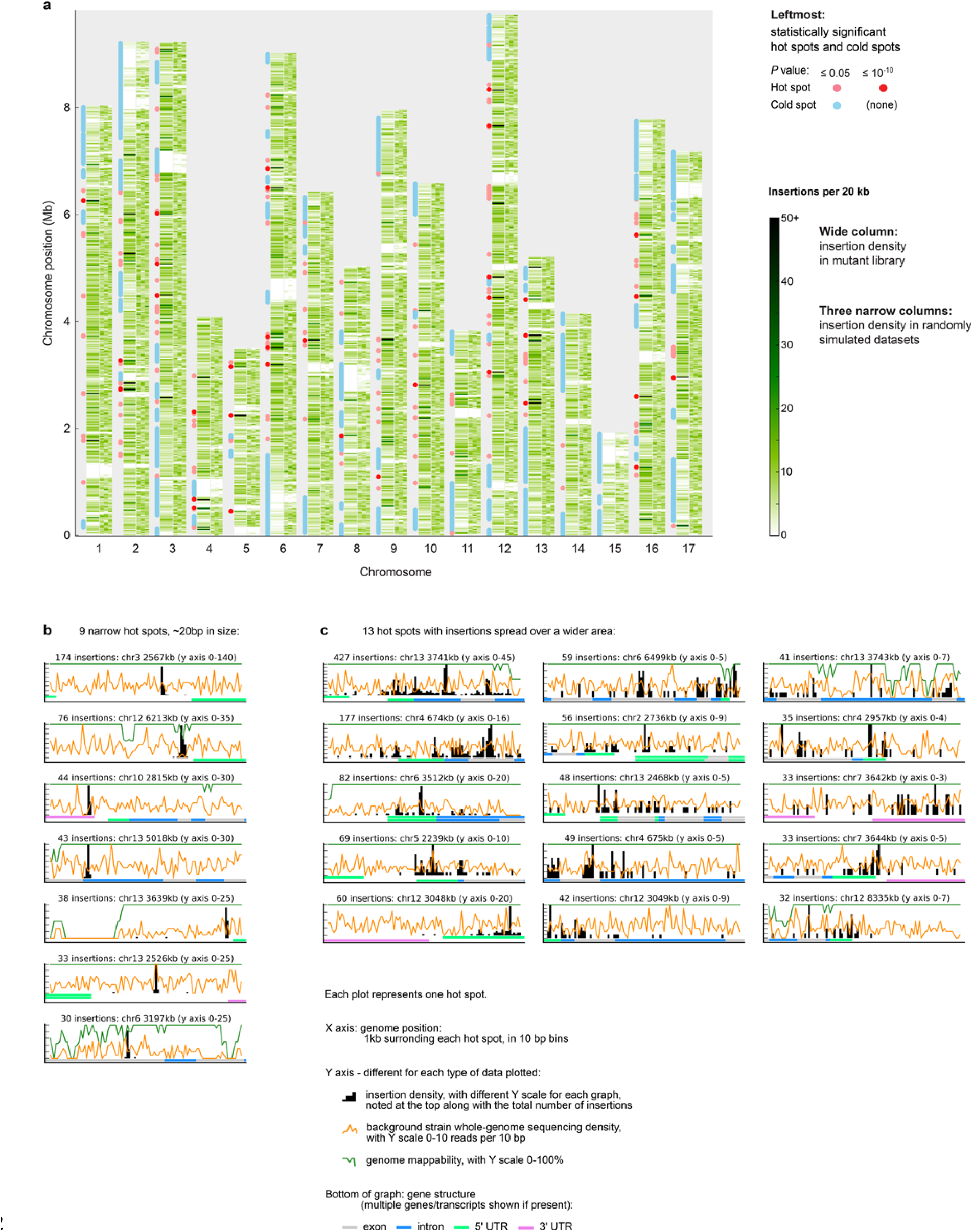
The distribution of insertions in the genome is largely random, and the hot spots fall into two classes. **a**, For each chromosome, the observed insertion density is shown as a heatmap in a wide column, followed by three narrow columns depicting three simulated datasets in which insertions were placed in randomly chosen mappable genomic locations. The simulated data provide a visual guide to the amount of variation expected from a random distribution. The large white areas present in both the observed and simulated data correspond to repetitive genomic regions in which insertions cannot be mapped uniquely. The red and blue circles/lines to the left of each chromosome show statistically significant insertion hot spots and cold spots, respectively. To ensure that we are showing true insertion density rather than artifacts caused by junk fragments or other mapping inaccuracies, the plot of insertion site distribution and identification of hot/cold spots are based on confidence level 1 insertions only. In contrast, Fig. 1c shows the distribution of insertions of all confidence levels over the genome. **b** and **c**, Each plot represents a 1-kb genomic region surrounding one hot spot, showing multiple features of that region, as listed in the legend. The plots shown are the 22 1-kb regions with the highest total insertion number. The total number of insertions for each region is listed above each plot, along with the genomic position and the y-axis range. **b**, 7 of the top 22 hot spots are narrow, with 20 or more insertions in a 10-bp area, and a total width of 20-30 bp with few or no additional insertions in the surrounding 1 kb. **c**, 15 of the top 22 hot spots are wider, with multiple peaks of high insertion density spanning at least hundreds of base pairs. In either class, the insertion density peaks do not appear to reliably correlate with any of the other genomic features shown. Data presented in this figure are described in the Supplementary Note.

**Supplementary Fig. 5.**
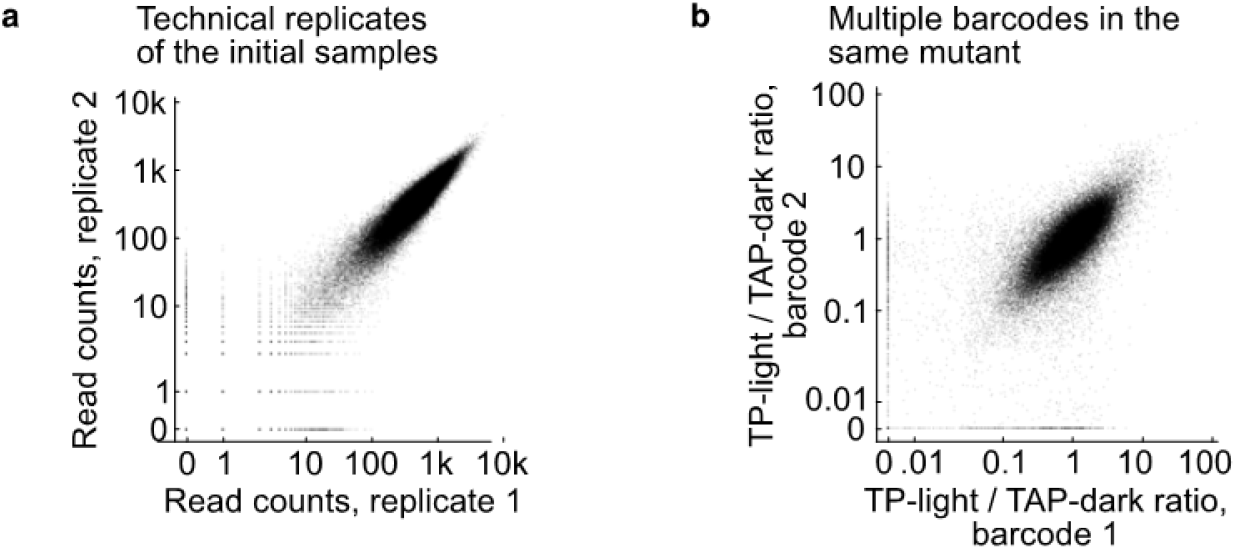
The barcode sequencing method is robust. **a**, The barcode sequencing read counts (normalized to 100 million total reads) for each insertion were highly reproducible between technical replicates, with a Spearman’s correlation of 0.978. 94% of barcodes showed a normalized read count of no more than a 2-fold difference between the two replicates. **b**, The TP-light/TAP-dark ratios of multiple barcodes in the same mutant are consistent, with a Spearman’s correlation of 0.744. Only 4% of insertion pairs had a greater than 5x difference between ratios. See also Fig. 3, b and c.

**Supplementary Fig. 6.**
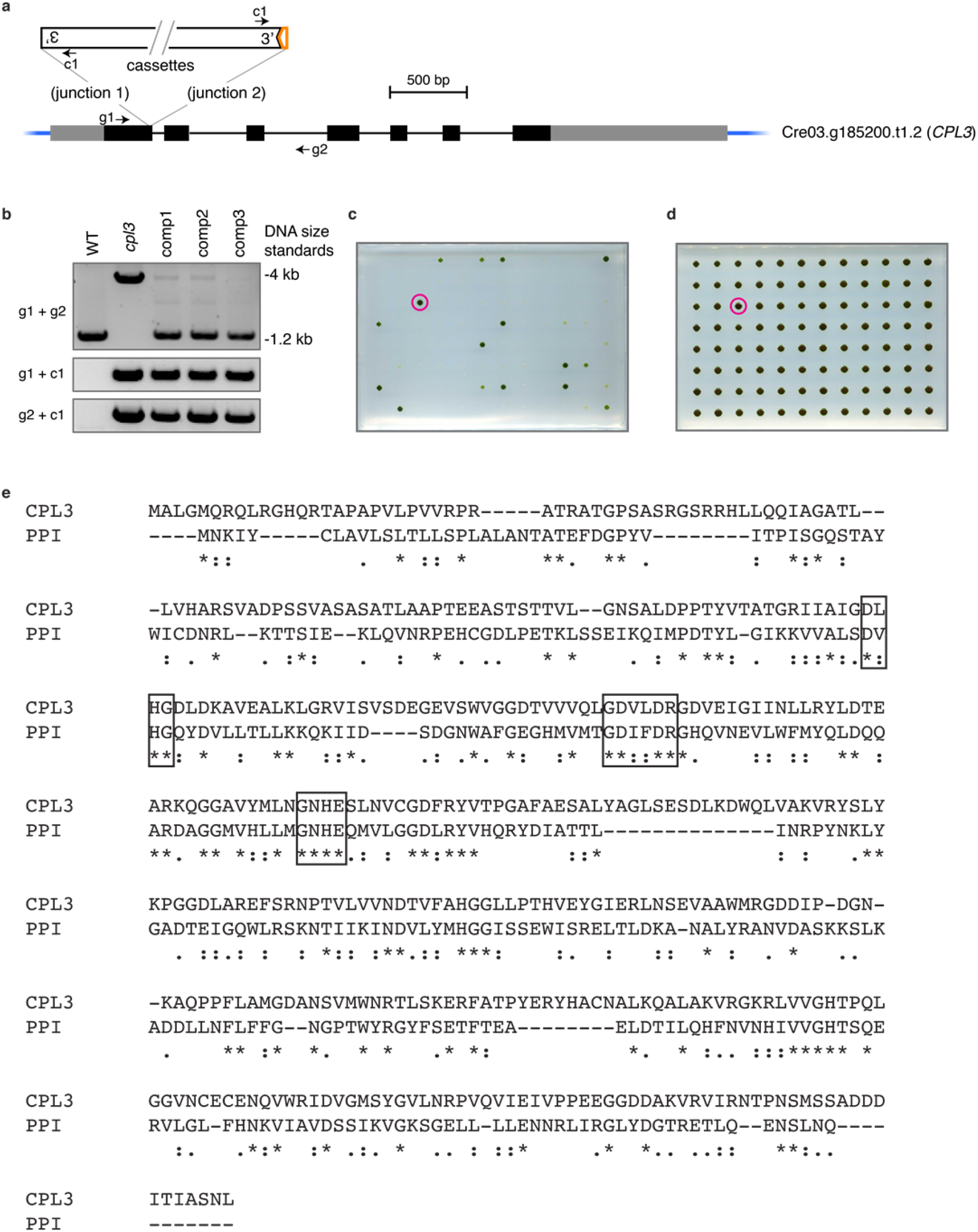
Molecular characterization of the *cpl3* mutant. **a**, The cassette insertion site is indicated on a model of the *CPL3* gene from the Chlamydomonas v5.5 genome. In the gene model, black boxes, gray boxes and thin lines indicate exons, UTRs, and introns respectively. Two cassettes are inserted in opposite orientations, with one of them truncated on the 3’ side (indicated by a notch); the 5’ ends may be intact or truncated. The orange box arrow indicates insertion of a small fragment of unknown origin. Binding sites for primers g1, g2, and c1 are indicated. **b**, PCR genotyping results of *cpl3* and complemented lines. PCR with the primer pair “g1 + g2” indicated presence of an insertion within the *CPL3* gene in the *cpl3* mutant and presence of wild-type *CPL3* sequence in the complemented lines. PCR with primer pairs “g1 + c1” and “g2 + c1” showed the presence of a cassette inserted into the *CPL3* gene in *cpl3* as well as the complemented lines. **c**, *cpl3* mutants transformed with the *CPL3* gene were arrayed and grown photosynthetically in the absence of acetate for one day under 100 µmol photons m^-2^ s^-1^ light and four additional days under 500 µmol photons m^-2^ s^-1^ light before imaging. The colony circled was a positive control strain that grows photosynthetically. Approximately 14% of transformants showed rescued photosynthetic growth, a frequency consistent with other genetic studies in Chlamydomonas^38^. **d**, The same transformants were grown for five days in the presence of acetate in the medium under 50 µmol photons m^-2^ s^-1^ light. All colonies grew similarly. **e**, CPL3 contains conserved tyrosine phosphatase motifs. Sequences of CPL3 in Chlamydomonas and its homolog psychrophilic phosphatase I (PPI) in *Shewanella sp*. were aligned using Clustal Omega^39^. Asterisks (*), colon (:), and period (.) indicate conserved, strongly similar, and weakly similar amino acid residues, respectively. The motifs that are conserved among multiple protein phosphatases^40^ are boxed. Data in panels **a-d** are described in the Supplementary Note. See also Fig. 4.

**Supplementary Fig. 7.**
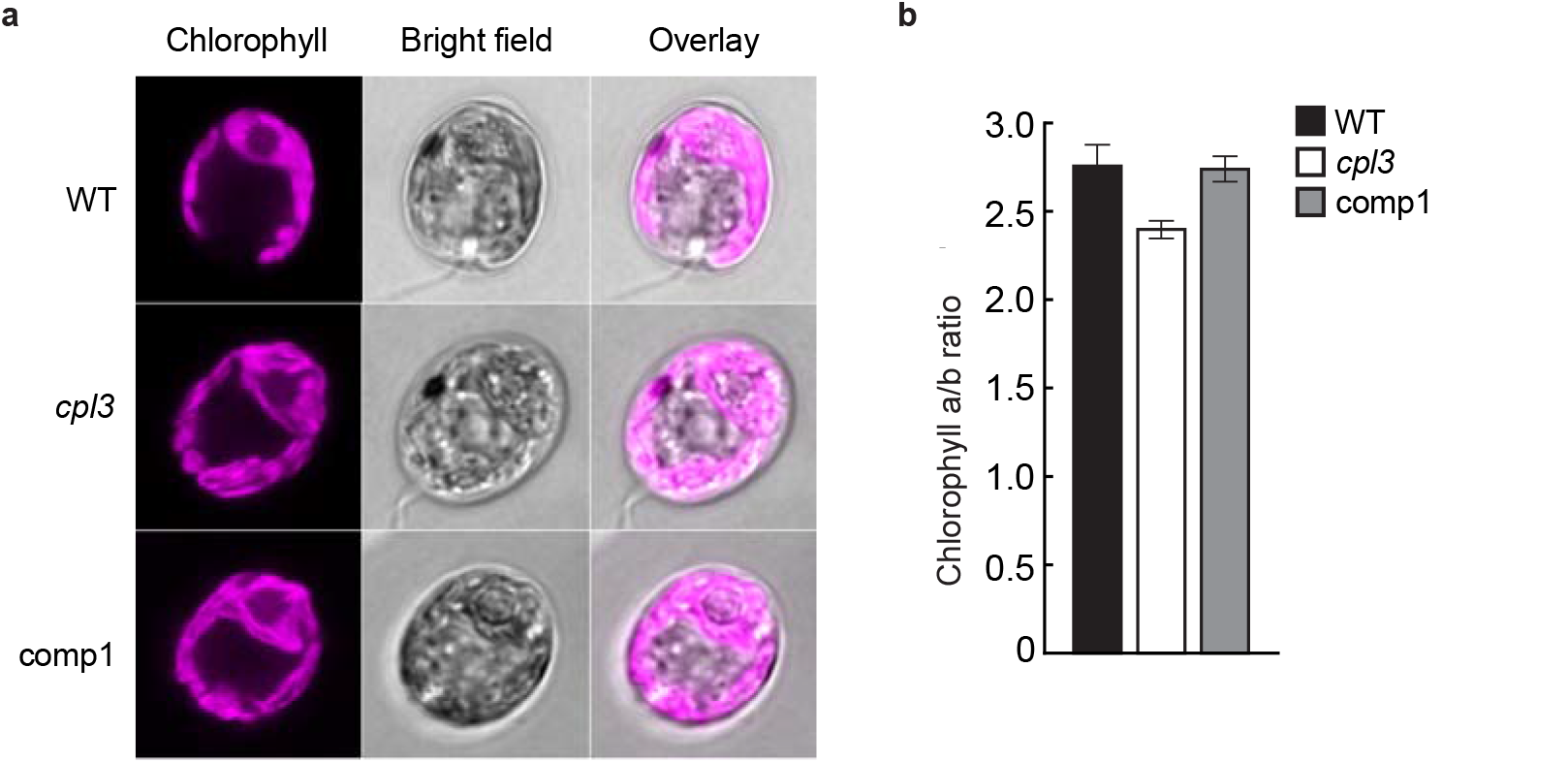
Phenotypic characterization of the *cpl3* mutant. **a**, *cpl3*, the wild-type strain (WT), as well as the complemented line (comp1), contain a normal cup-shaped chloroplast. Representative images of confocal chlorophyll fluorescence, bright field, and an overlay are shown for each strain. **b**, *cpl3* has a lower chlorophyll *a*/*b* ratio than WT and comp1 (*P*<0.03, Student’s *t*-test).

### Captions for Supplementary Table 1 to S14 (each provided as a separate file)

**Supplementary Table 1 | Primers and experimental design for all PCRs related to library generation and mapping**

This database includes primers used for generation of the insertion cassette (Supplementary Fig. 1a), primers used for barcode amplification and sequencing (Fig. 3a, Supplementary Fig. 1f, and Supplementary Fig. 2a, b), primers and experimental design for LEAP-Seq (Supplementary Fig. 1g and Supplementary Fig. 2f-j). See also Methods.

**Supplementary Table 2 | Binary codes for plate super-pooling**

In this database, each of the 570 rows corresponds to a plate-pool and each of the 21 columns corresponds to a plate-super-pool. A 1 in row X and column Y indicates that plate-pool X was included in plate-super-pool Y, and a 0 indicates that it was not. See also Supplementary Fig. 1f, Supplementary Fig. 2a, and Methods.

**Supplementary Table 3 | Binary codes for colony super-pooling**

Binary codes for the generation of 20 colony-super-pools from 384 colony-pools are shown in the same format as in Supplementary Table 2. See also Supplementary Fig.1f, Supplementary Fig. 2a, and Methods.

**Supplementary Table 4 | Read counts for each barcode in each combinatorial super-pool**

The columns in this database are barcode, side (5’ or 3’), and read counts for that barcode in each super-pool, normalized to 1 million total reads for each side and each super-pool.

Only barcodes deconvolved to both a plate and a colony are included. See also Supplementary Fig. 1f, Supplementary Fig. 2a and Methods.

**Supplementary Table 5 | List of all mapped mutants in the library**

Each line is an insertion junction (i.e. one mapped side of an insertion). Some insertions have one mapped side, some have two. Some mutants have multiple insertions. The columns are as follows (explained in detail in Methods):

- mutant_ID - the ID of the mutant if it was included as part of the “consolidated” set for long-term maintenance, ‘-’ otherwise. A mutant can have multiple insertions.
- side - which side of the cassette the data is derived from, 5’ or 3’. · chromosome, strand, min_position - the mapped position of the insertion, potentially corrected for a junk fragment, depending on the value of the if_fixed_position column.
- gene, orientation, feature, gene_end_distances - the gene containing the insertion, the orientation with respect to the gene, the feature of the gene, and the distance from the 5’ and 3’ end of the gene. If the position is inside two overlapping genes, all fields will have two entries separated by ‘&’. For intergenic positions, all values are ‘-’.
- intergenic_adjacent_genes, intergenic_orientations, intergenic_gene_distances – for intergenic positions, these fields note the two adjacent genes, the position of the insertion with respect to those two genes, and the distance from them. Each field will have two entries separated by ‘&’, unless the insertion position is on the edge of a chromosome and has no gene on one side. For insertions in genes, all values are ‘-’.
- if_both_sides – ‘-’ if the insertion only has one mapped side, otherwise ‘perfect’, ‘deletion’ or ‘duplication’ for insertion junctions that are two sides of a confidence level 1 insertion depending on whether there was a deletion/duplication in the genomic DNA, or ‘with-junk’ for insertion junctions that are two sides of a confidence level 2 insertion with a junk fragment on one side.
- confidence_level - the confidence level for the insertion mapping, as described in Supplementary Fig. 3a.
- if_fixed_position – ‘no’ if no junk fragment was detected in the insertion (confidence level 1, 3, the side of confidence level 2 with no junk fragment, and a small fraction of confidence level 4); ‘yes_nearest_distal’ if a junk fragment was detected and corrected. For the junk fragment sides of confidence level 2 insertions, the value is just ‘yes_nearest_distal’, indicating that the corrected insertion position for this line is the position of the nearest distal LEAP-Seq read; for most confidence level 4 insertions, the value is ‘yes_nearest_distal_+/-X’, indicating a further correction of X bp that was applied to the position to compensate for the average distance between the nearest distal LEAP-Seq read and the true insertion position (see Methods).
- LEAPseq_distance, LEAPseq_percent_confirming - the highest distance spanned by a proximal and distal LEAP-Seq read pair mapping to the same region, and the fraction of pairs that map to the same region (see Supplementary Fig. 1g and Supplementary Fig. 2f-j).
- flanking_seq - the flanking sequence immediately adjacent to the cassette, or, if the if_fixed_position column value is not ‘no’, the sequence of the distal LEAP-Seq read closest to the corrected mapping position.
- barcode - the barcode sequence of the insertion.
- gene_name, defline, description, etc - gene annotation from Phytozome^10^.

**Supplementary Table 6 | Primers and results of PCRs used to verify the insertion sites of randomly-picked mutants from the mutant library**

For column definitions, see the legend of Supplementary Table 5.

**Supplementary Table 7 | Statistically significant insertion hot spots and cold spots**

The columns give the hot spot position (chromosome, start and end base number), type (hot spot or cold spot, i.e. enriched or depleted in insertions), false discovery rate (FDR), number of insertions in the spot, and the expected number of insertions based on the length and mappability of the spot (see Methods). Only hot spots that passed the filtering are listed.

**Supplementary Table 8 | Statistically significant depleted functional terms**

The columns give the gene ontology (GO) term, FDR, the ratio of observed vs expected insertions in all the genes annotated with the term (i.e. the effect size), the number of genes annotated with the term, the total number of insertions in those genes, the total mappable length of those genes, and the GO term definition (see Methods). Only depleted GO terms are listed - most of the enriched GO terms were due to hot spots in a single gene.

**Supplementary Table 9 | Candidate essential genes**

The columns give the gene ID, its mappable length (excluding UTRs), the number of insertions in the gene (again excluding UTR insertions), the expected number of insertions given the mappable length if the insertion distribution was random, the FDR, and gene annotation from Phytozome^10^ (data described in the Supplementary Note).

**Supplementary Table 10 | Read counts of barcodes before and after pooled growth in the photosynthesis screen**

The columns give the barcode, the gene in which the insertion is located (or “-” if intergenic), the gene feature, the side of cassette the data is derived from, and deep-sequencing read numbers (raw and normalized to 100 million) in the two technical replicates of the initial pool, the pool after growth in TAP-dark, and two biological replicates after growth in TP-light.

**Supplementary Table 11 | Statistics of the pooled growth data for all genes**

The columns give:

- the gene ID and name, the hit_category (higher-confidence candidate or lower-confidence candidate, otherwise “-”).
- the number of alleles in the gene with and without a phenotype in replicate 1, the resulting *P* value and FDR in replicate 1 (see Methods; genes with only 1 allele have no FDR).
- the same four numbers for replicate 2.
- PredAlgo-predicted localization for the gene: C = chloroplast, M =mitochondrion, SP = secretory pathway, O = other, and “-” if no prediction could be made.
- gene annotation data from Phytozome^10^.

**Supplementary Table 12 | Summary of previous characterizations of higher- and lower-confidence genes’ roles in photosynthesis**

The columns are similar to those in Supplementary Table 11, but additionally include:

- Previously reported function in photosynthesis for each gene.
- The corresponding references.

**Supplementary Table 13 | Read counts of *cpl3* exon and intron alleles in the pooled screens**

The numbers of read for each barcode per 100 million total reads are presented. Some of the mutants contain two barcodes and are labeled with an asterisk (*). In such cases, the barcode with a greater TAP-dark read count is used to determine whether there is a phenotype. The mutants that fall below the phenotype cutoff are labeled with an obelisk (†). Two out of the seven mutants were included in the screen but not included in the consolidated set.

**Supplementary Table 14 | Proteomic characterization of the *cpl3* mutant**

The columns give:

- ID for nuclear/chloroplast-encoded genes.
- name - gene name from Phytozome bulk annotation.
- annotations for proteins plotted in Fig. 4 - names of the complexes and protein subunits are shown.
- spectral counts – number of spectra detected for peptides derived from each protein, related to protein abundance.
- WT_repl1/2/3, *cpl3*_repl1/2/3, comp1_repl1/2/3 - log_2_ intensity values for three replicates of each sample, normalized to a median of 0. WT, the wild-type parental strain CC-4533. Comp1, the complemented line.
- WT_mean, *cpl3*_mean, comp1_mean - the average of the three replicate intensity values for each sample.
- WT-*cpl3*_diff - *cpl3*_mean subtracted by WT_mean. The lower this value, the less abundant the protein is in *cpl3* relative to WT.
- WT-*cpl3*_pval - the raw *P* value comparing the normalized replicate intensities for WT and cpl3, using an unpaired t-test.
- WT-*cpl3*_FDR - the false discovery rate (i.e. the *P* value adjusted for multiple testing, using the Benjamini-Hochberg method^12^).
- WT-comp1_diff, WT-comp1_pval, WT-comp1_FDR, *cpl3*-comp1_diff, *cpl3-*comp1_pval, *cpl3*-comp1_FDR - the same three values for the comparison between WT and comp1 samples and between *cpl3* and comp1 samples.
- defline, description, and all further columns - gene annotation from bulk Phytozome data.

